# Including organism and environmental heterogeneity in kinematic continuum models of collective behaviour with applications to locust foraging and group structure

**DOI:** 10.1101/2023.08.29.555245

**Authors:** F. Georgiou, J. Buhl, J.E.F. Green, B Lamichhane, N. Thamwattana

## Abstract

Collective behaviour occurs at all levels of the natural world, from cells joining together to form complex structures, to locusts interacting to form large and destructive plagues. These complex behaviours arise from simple individual and environmental interactions, and thus lend themselves well to mathematical modelling. One simplifying assumption, that of relative homogeneity of organisms, is often applied to keep the mathematics tractable. However, heterogeneity arising due to the internal state of individuals has an impact on these interactions and thus plays a role in group structure and dynamics.

In this paper, we introduce a continuum model that accounts for this heterogeneity in the form of a state space that models an organisms internal state and converts this to movement characteristics. Using a variety analytic techniques, we investigate the effect of internal state on aggregation density and aggregation formation behaviour, finding that density and formation is most affected by the ratio of attractive to dispersive interactions.

We then apply the model to a concrete example of locust foraging to investigate the effect of food, hunger, and gregarisation on locust group formation and structure. Through numerical simulations we find that the most gregarious and satiated locusts tend to be located towards the centre of locust groups. Conversely, hunger drives locusts towards the edges of the group. Finally, we find that locust group dispersal may be driven in part by hunger.

**Author Summary:** Collective behaviour occurs at all levels of the natural world, from cells joining together to form complex structures, to locusts interacting to form large and destructive plagues. We can modelling these complex behaviours using mathematical models however we often need to rely on simplifying assumptions to keep the mathematics easy enough to analyse. One simplifying assumption that is often employed is assuming that all the modelled organisms are the same (or in one of only a few possible states). However, this is often not the case in nature where the differences between individuals arising due to internal characteristics, such as hunger or age, often affects their behaviour and thus can change group dynamics.

In this paper we introduce a mathematical model that is able to capture these differences and apply the newly developed model to locust foraging. We find that hunger tends to drive individuals to the edges of aggregations as well as lowers the maximum possible density. These two results combine to give a possible mechanism for the dispersal of locust groups.

## 1 Introduction

Collective behaviour occurs at all levels of the natural world, from cells joining together to form complex structures [42], to locusts interacting to form large and destructive plagues [46]. Collective behaviour can be defined as emergent and self-organising macroscopic collective motion and pattern formation arising from simple inter-individual and environmental interactions [6, 48, 49]. Due to the simple nature of these interactions, mathematical models have proven to be an important tool in studying this phenomenon [4]. These mathematical models can be broadly classified as either discrete, where organisms are modelled as individuals or continuum in which they are represented as a continuous density function [4].

Self propelled particle (SPP) models are a frequently used discrete approach in which organisms are modelled as individual points which update their velocity according to simple interaction rules. SPPs can be further categorised as second order models if they include particle inertia, and first order (or kinematic) if inertia is neglected [16]. Second order SPP models have been used fairly extensively as they are able to capture collective movement mechanisms such as alignment or pursuit/escape interactions [3, 11, 12, 13]. First order SPP models, where drag is assumed to dominate over inertia have been applied for modelling stationary pattern formation, such as cellular organisation [17], or disordered group behaviour such as the rolling locust swarm [7, 44].

One downside of discrete models is that there are few analytical tools available to study their behaviour. In contrast, continuum models, in which organisms are represented as a population density that is a function of space and time, can be analysed using an array of tools from the theory of partial differential equations (PDEs). They are most appropriately employed when there are a large number of individuals since they do not account for individual behaviour, instead giving a representation of the average behaviour of the group. The latter (continuum) approach is adopted in this paper.

The non-local aggregation equation, first proposed by Mogilner and Edelstein-Keshet to model swarming behaviour [31], is a common continuum PDE analogue of the kinematic SPP model [9, 10]. It consists of a conservation of mass equation for the population density *ρ*(*x, t*) given by

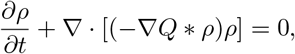

where *Q* is defined as some social interaction potential and *\** is the convolution operation. For this type of model, the existence and stability of swarms has been proven, and both travelling wave solutions [31] and analytic expressions for the steady states [7] have been found. This model has been further extended to include non-linear local repulsion which leads to compact and bounded solutions [43]. Usually used for single populations, the model has been further adapted to consider multiple interacting populations [23].

One key consideration is that in the interest of mathematical tractability continuous models typically assume a homogeneity amongst organisms. That is, individuals are indistinguishable from each other or only vary in one or two discrete ways, e.g. solitarious/gregarious locusts [22, 45]. In real world animal social grouping, phenotypic and environmental heterogeneity plays an important role in group structure and dynamics [27]. The phenotypic heterogeneity arises due to slow changing attributes such as age, size, personality, and/or fast-changing factors such as energy reserves. For example, hungrier individuals tend to migrate towards the front of moving groups in both caterpillars [30] and crimson spotted rainbow-fish [25], while faster pigeons hold a leadership position in flocks [35]. Whilst numerous examples of modelling heterogeneity exist, they are however dominated by SPP models [4], with very few examples of continuous ones [32], and to our knowledge none that include both phenotypic and environmental heterogeneity. In this paper, we derive a kinematic model for collective behaviour which considers both phenotypic heterogeneity (in the form of an internal state) and environmental interactions. We then explore the model using a the locust as a prototypical example.

Locusts are short horned grasshoppers that exhibit density-dependent phase-polyphenism, i.e., two or more distinct phenotype expressions from a single genotype depending on local population density [34]. In locusts there are two key distinct phenotypes, solitarious and gregarious, with the process of transition from solitarious to gregarious called gregarisation (and the reverse, solitarisation). Gregarisation affects many aspects of locust biology from colouration [38], to reproductive features [39], to behaviour [40]. Behaviourally, solitarious locusts are characterised by an active avoidance of other locusts, whereas gregarious locusts are strongly attracted to other locusts. Gregarisation is brought about by locusts crowding together and can be reversed by isolating the individuals [47]. In the Desert locust (*Schistocerca gregaria*), gregarisation can take as little as 4 hours with the time-frame for reversal dependant on the length of time the individual has been gregarious (again, potentially as little as 4 hours) [34].

It is in this gregarious state that adult locusts form the infamous large and destructive plagues. Prior to this, juvenile locusts form hopper bands, large groups of individuals (can be in the millions) marching in unison [47]. Earlier continuum models of locusts have attempted to capture the onset of collective behaviour by modelling the phase polyphenism as two distinct sub-populations, with some density dependent transition between the two [22, 45]. However, this relationship might be more continuous than captured by that assumption [2]. In addition, models assumed outside of locust-locust interactions other behavioural aspects are constant, however this is not the case as their movement characteristics are dependent on a variety of factors. For example, locusts modify their dispersal behaviour based on hunger [18, 36, 41]. It is these interactions of internal state that we aim to understand.

In this paper, we first present our model in Section 2 (and Appendix A), before obtaining analytic results of maximum density and the conditions required for the formation of aggregations in Section 3. We then look, in Section 4, at applying the model to locust foraging and investigate the effect of a continuous model of gregarisation and hunger on group formation and structure. Finally, we discuss our results and further avenues of exploration in Section 5.

## 2 Model

We begin by presenting a continuous kinematic model of collective behaviour that includes both local and non-local inter-individual interactions, as well as environmental interactions, with all interactions mediated by the internal state of the organism. We make the following modelling assumptions;

1. Organisms can be classified by their internal state.
2. Environmental interactions are local in nature.
3. Local interactions between organisms are repulsive (i.e. they avoid close physical contact).
4. Organisms experience a non-local (i.e. longer-ranged) interaction with organisms in any state.
5. The nature of all interactions are mediated by the organisms internal state and local environmental conditions.

In this model organisms are represented as a density of individuals (number per unit area) at point ***x*** in space, at time *t*, and with a *N* -dimensional internal state, ***n*** = (*n*_1_, .., *n*_*N*_), where each state can take a value on the interval [0, 1]. We will term the combination of all possible internal states as the state space (i.e. the state space is an *N* -dimensional hypercube). This state may characterise their satiation, age, stage of gregarisation, etc. as a fraction of the greatest possible value, i.e. all the states are normalised and unit-less. Let the density of organisms in space, state, and time be given by, *ρ*(***x, n***, *t*) with the total local density in space defined as

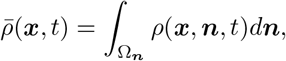

where Ω_***n***_ is our complete state domain, we will also define Ω_***x***_ as our spatial domain. For later convenience we will also define the total mass of organisms as

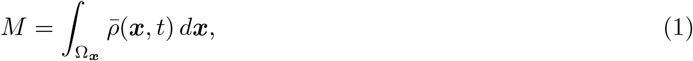

as well as a generic term for the environment, *E*, that might represent such environmental conditions as food density, sunlight, temperature, etc.

We assume that the time-scales we are investigating are shorter than the life cycle of the organism, ignoring births and deaths and thus conserving the total number of organisms. We consider local organism-organism interactions (e.g. crowding), local organism-environment interactions (e.g. foraging), and non-local organism-organism interactions (e.g. sight and smell mediated interactions between individuals). We also allow for a movement within the state space (e.g. the change in an organisms hunger level). Hence, conservation laws give equations of the form

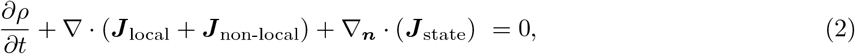

where ***J*** _local_ is the flux due to local interactions, ***J*** _non-local_ is the flux due to non-local interactions, ***J*** _state_ is the flux around the state space and *∇*_***n***_ represents the differential operator, *∇*, applied to the state space variables, i.e.

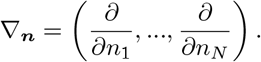

For our local interactions we follow the work of Painter and Sherratt [33] and derive them as the limit of a lattice model (for the full derivation see Appendix A). By assuming that organism movement due to local interactions depends on local environmental conditions, the state of the organism, and local population density. These assumptions gives our local flux as,

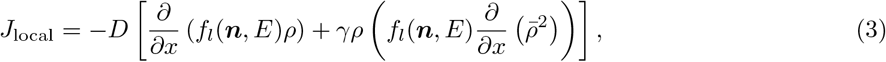

where, *f*_*l*_(***n***, *E*) relates state and local environmental conditions to movement, *D* is the linear diffusion coefficient, and *γ* is the non-linear diffusion coefficient.

Next, for our non-local interactions, we adopt the fluxes used originally by Mogilner and Edelstein-Keshet [31], and in numerous subsequent studies [22, 43, 45]. However, now we assume that the social potential is only a function of space (i.e. its constant in state and time) and the strength (and direction) is mediated by both the internal state and the environment, giving

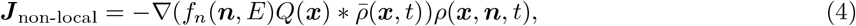

where *Q*(***x***) is the social potential and *f*_*n*_(***n***, *E*) is a function relating internal state and environmental conditions to the strength and direction of the non-local force.

Finally, we assume that organisms change their internal state based on some *N* dimensional velocity vector, ***v***_*n*_ giving our flux as

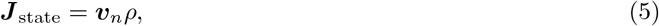

with each element of 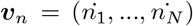 being a function describing the rate of change in state. Specific examples will be derived in Section 4.

Combining (3), (4), and (5) into (2) we get

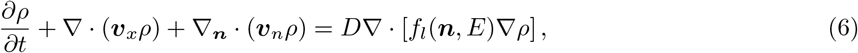

with

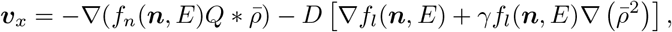

For the analytic results we try to keep to the most general equations possible. However, when a concrete example is required, we use the common social potential given by

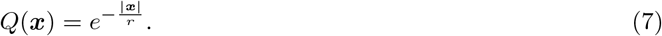

For boundary and initial conditions, they are presented in the relevant sections. Finally, we have not given evolution equations for the environment, *E*, as this depends on the modelled situation, we give an example later in Section 4.

## 3 Analytic Results

We now gain some insight into the models behaviour through PDE analysis techniques. We begin by using gradient flow methods [1] to look at the maximum density of aggregations with environmental conditions that are constant in both space and time, as well as homogeneous organism state. We then use linear stability analysis to look at the conditions required for aggregations to form with all organisms being in the same state, then in one of two states. While we have in essence assumed away the heterogeneity of our model organism and environment, the results found here can give insights into the behaviour under more heterogeneous conditions. Finally, the complete calculations can be found in Appendix B.

### 3.1 Density of aggregations

In order to facilitate analysis, we begin by introducing some simplifying assumptions, under which we can estimate the maximum density and width of a aggregations at both the large and small mass limits in one dimension (i.e. as *M → ∞* and *M →* 0, respectively). To begin, our assumptions are the environment, *E*, is constant in space and time, and all the organisms are in the same unchanging state i.e. 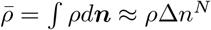 where *N* is the number of state dimensions and Δ*n* represents a small area in state space. Finally, we will label the support of *ρ* as Ω^*′*^. These assumptions give a gradient flow of the form

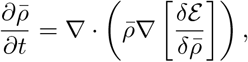

where

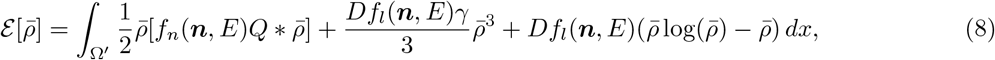

with the minimisers satisfying

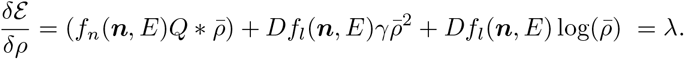

Full details of the calculations can be found in Appendix B.1.

#### 3.1.1 Large mass limit

We begin by investigating the maximum density and support as *M → ∞*. Taking (8) and then further assume the following: 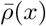 is approximately rectangular. Additionally, while the support of 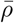, Ω^*′*^, is infinite due to the linear diffusion the bulk of the mass is contained as a series of aggregations, we will approximate the support of an aggregation as Ω. Finally, for a single aggregation we assume that the support is far larger than the range of *Q*. We thus approximate *Q ≈ V*_*Q*_*δ*(*x*), where *δ*(*x*) is the Dirac delta function, with *V*_*Q*_ = *∫ Qd****x***. We can then find our maximum density, 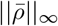, and aggregation support, ∥Ω∥, as

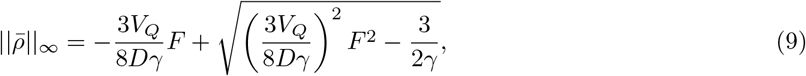

with support

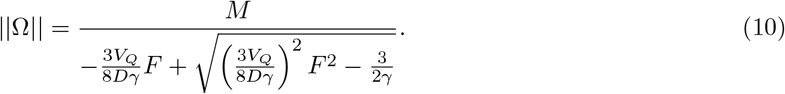

where

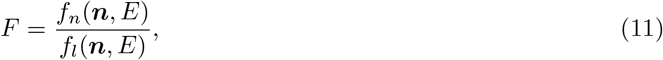

is the ratio of our non-local and local forces. There are a few things to note in these equations. Firstly for aggregations to exist, i.e. 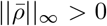, *F* must be less than 0 corresponding to an attractive social potential (or an attractive local movement; however, if both local and non-local components are attractive our original assumption of 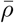 being rectangular would not hold as the equation is singular). Next, from this we find that any change in state, ***n***, that increases the rate of local movement without a corresponding increase in non-local movement would decrease the maximum density of aggregations and thus increase the size of the support (and vice versa). Finally, any change in state that increases the rate of non-local movement compared to local movement would increase the maximum density of aggregations and thus decrease the size of the support.

We can also use (9) to estimate the parameter *γ*, given a maximum density of organisms, *ρ*_*∞*_, we find,

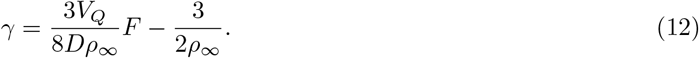

#### 3.1.2 Small mass limit

We now turn to the case as *M →* 0. Assuming that for a single aggregation we can approximate the social interaction potential using a Taylor expansion. In this section we use *Q*(*x*) given by (7), with 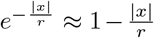.Additionally, we ignore the effect of linear diffusion within Ω (to make the calculations possible), giving our maximum density, 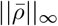, and aggregation support, ∥Ω∥, as

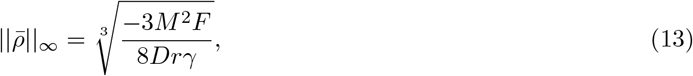

and

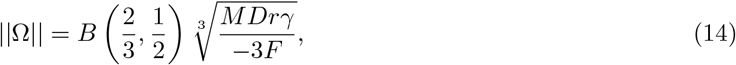

which gives a similar relationship as the large mass limit however here both the maximum density and support scale with mass, *M*.

#### 3.1.3 Comparison of mass limits and simulations

We can check the accuracy of our estimates by comparing them to simulation results (Details of the numerical scheme can be found in Appendix C [15, 19, 26, 28]). We begin by defining two dimensional state space, (*n*_1_, *n*_2_), with the first dimension only affecting the non-local component of force and the second dimension affecting the local component. For example, these could be considered in locusts to represent gregarisation and hunger, respectively. We first let,

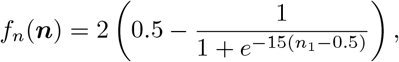

and

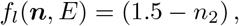

in addition we let,

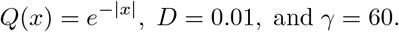

For each simulation we place a mass of organisms, *M*, in every combination of the states *n*_1_ = 0.85 (results for *n*_1_ = 0.75, 0.85 and 0.95 can be seen in Appendix B) and *n*_2_ = 0.15,0.55,0.95 in the centre of the domain (we vary the domain size according to mass so there is no boundary interaction) and run it to a pseudo steady state, *t* = 1000.

The results for 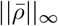 and ∥Ω∥ can be seen in Figure 1. In the plots, the dotted lines represent the minimum estimates between the small and large mass limits, and the solid lines represent simulated results. As ∥Ω∥ is theoretically infinite due to the linear diffusion, for the simulated ∥Ω∥ we select the region for which 98% of the mass, *M*, is contained. We can see that as *M* increases the simulated limits approach those given by the theoretical estimates, however there is some error likely due to error in the simulations and our method of approximating ∥Ω∥. In addition, the small mass estimates are considerably less accurate than the large mass limits (due to ignoring linear diffusion), however, the estimates display qualitatively similar behaviour.

**Figure 1.**
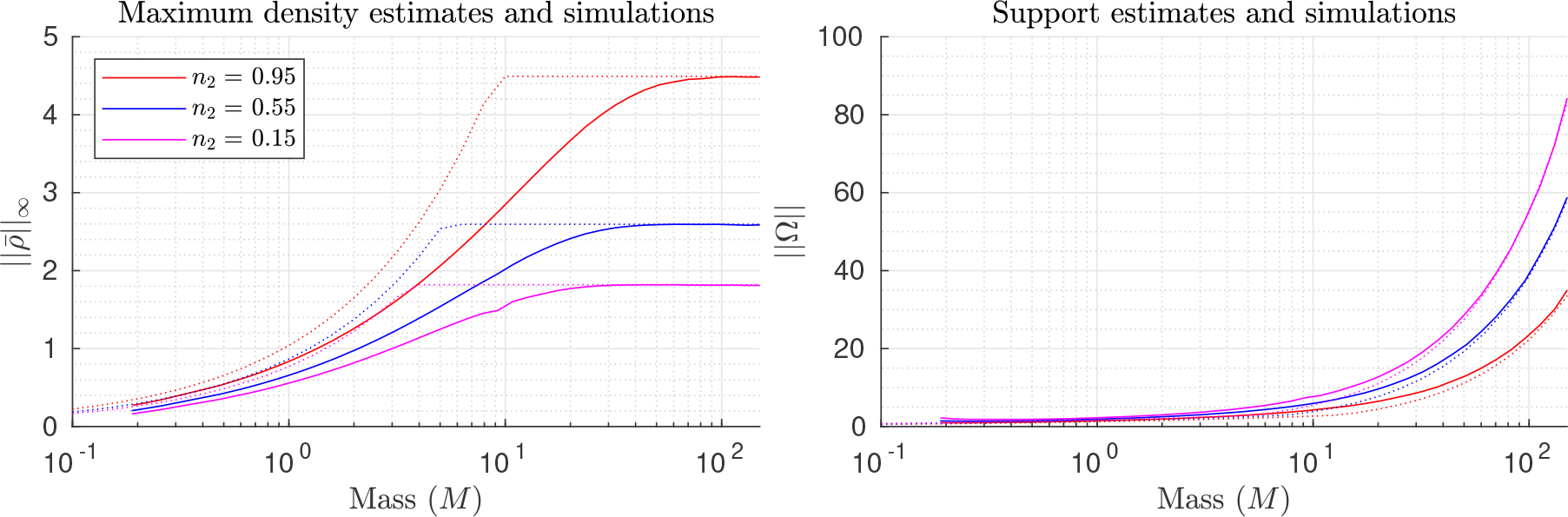
Small and large mass max density and support, estimates and simulations. Estimates and simulations for the maximum density (left) ans size of the support (right). Estimates for different states are plotted using dotted lines, with the corresponding simulation results plotted in solid lines. As ∥Ω∥ is theoretically infinite due to the linear diffusion, for the simulated ∥Ω∥ we select the region for which 98% of the mass, *M*, is contained.

We can delve deeper into the estimates by looking at individual simulation result. In Figure 2, we see the small (*M* = 0.48) and large *M* = 96.17 mass limit estimates (left and right plots respectively) with their corresponding simulations for *n*_1_ = 0.95, *n*_2_ = 0.15, 0.55, 0.95. In the plots, the vertical dotted lines are the estimates of the support, the horizontal dotted lines are the estimates of the maximum density, and the solid lines are the simulation results. We can see that the large mass estimates are considerably more accurate but the small mass estimates do capture some of the qualitative behaviours.

**Figure 2.**
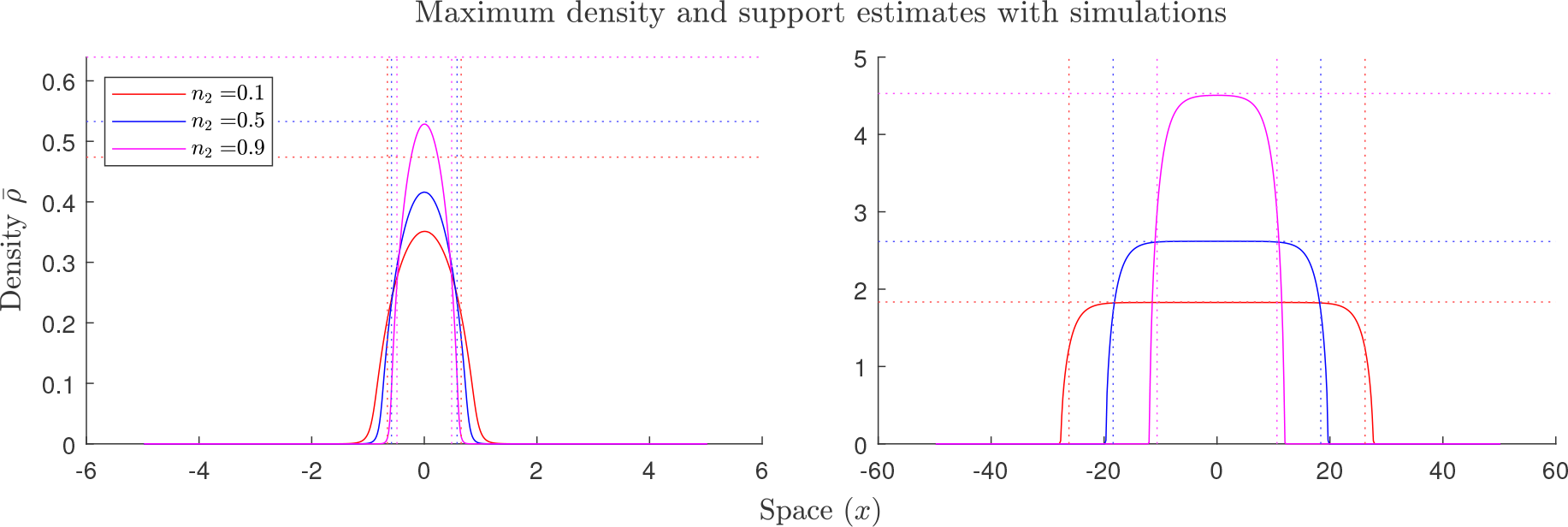
Small and large mass support estimates and simulations. Simulations of small (*M* = 0.48) and large *M* = 96.17 mass limit estimates (left and right plots respectively) with *n*_1_ = 0.95, and *n*_2_ = 0.15, 0.55, 0.95. In the plots, the vertical dotted lines are the estimates of the support, the horizontal dotted lines are the estimates of the maximum density, and the solid lines are the simulation results.

### 3.2 Linear stability analysis of homogeneous steady states

In order to gain insights into the conditions under which aggregations can form, we investigate the stability of spatially-homogeneous steady states. In this analysis we perturb the homogeneous steady state by adding a small amount of noise. We then find under what conditions (given in the form of an inequality) the small perturbations grow and are likely to lead to aggregations. We again assume that *E* is constant in space and time. Full details of the calculations can be found in Appendix B.2.

#### 3.2.1 A single state

We begin by considering all organisms in a fixed single state, ***n***_*f*_, i.e. 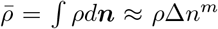 where *m* is the number of state dimensions. These assumptions give our condition for instability in terms of our two state based force, as

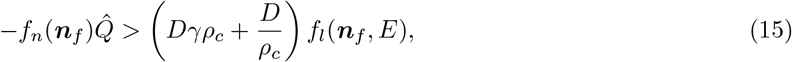

or alternatively

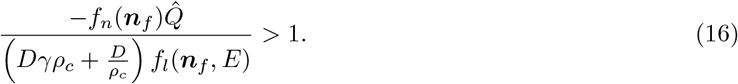

From this we can see that if *f*_*n*_(***n***_*f*_) *>* 0, corresponding to a repulsive non-local term, then aggregations will not form and if *f*_*n*_(***n***_*f*_) *<* 0 then non-local forces simply need to be greater than local forces for aggregations to form.

#### 3.2.2 Two states

We can also consider two distinct sub-populations in states ***n***_*s*_ and ***n***_*g*_ with *f*_*n*_(***n***_*g*_, *E*) *< f*_*n*_(***n***_*s*_, *E*) (i.e. *f*_*n*_(***n***_*g*_, *E*) is less repulsive than *f*_*n*_(***n***_*s*_, *E*), or even attractive). The fraction of the population in state ***n***_*g*_ is given by *ϕ*_*g*_. This allows us to rephrase (6) as a system of PDEs given by,

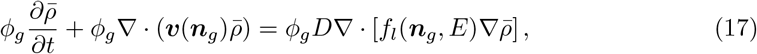

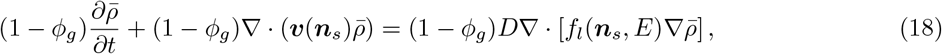

with

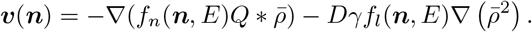

We then find the condition for instability in terms of *ϕ*_*g*_ as

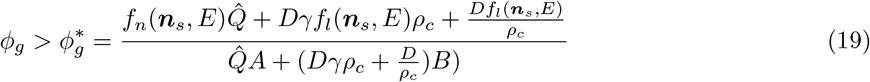

where

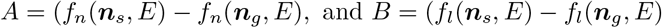

From this we can see that: in environmental conditions that reduce dispersal the gregarious fraction required for aggregation formation decreases. The numerator is only in terms of ***n***_*s*_, thus states that increase the local and non-local terms will increase the gregarious mass fraction required for aggregation formation. The denominator depends on the difference between the states: Locally, if the gregarious state has less local movement than the solitarious state this decreases the gregarious mass fraction required for aggregation formation and vice-versa. Non-locally, decreasing the non-local force of both the gregarious and solitarious states decreases the gregarious mass fraction required for aggregation formation and vice-versa (as *f*_*n*_ *<* 0 is attraction and *f*_*n*_(***n***_*g*_, *E*) *< f*_*n*_(***n***_*s*_, *E*)). Finally, as 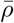 increases the gregarious fraction required for aggregation formation increases suggesting an upper organism density in order to transition away from the homogeneous steady state.

For our specific function 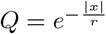, we get,

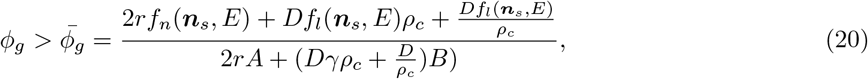

where

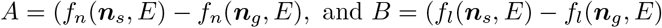

From this we can find the maximum homogeneous density, *ρ*_*c*_, that aggregations can still form. So taking (20) and substituting *ϕ*_*g*_ = 1 gives a maximum homogeneous density for aggregations as,

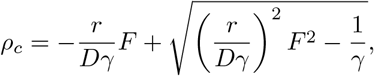

where *F* is given by (11) for the state ***n***_*g*_. Rather neatly, if 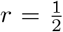, corresponding to 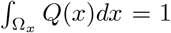, this becomes

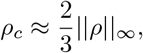

where ∥*ρ*∥_*∞*_ is given by (9), this is similar to the relationship previously derived by us [22].

## 4 Applications to locust foraging

In this section we use the model to investigate the effect of food and different state combinations on group formation and group heterogeneity within the context of locust foraging. We begin by stating our environmental variables and functions before deriving state dimensions and parameters of gregarisation and hunger. Then, we look at the derived functions and their effect on group formation and density based on the results form Section 3. Finally we perform a series of numerical experiments that builds upon our previous work in which locusts were modelled with a binary solitarious/gregarious state and food was considered without hunger [22].

### 4.1 Environment and state dimensions

Here we describe, derive and analyse an application of our model with two state dimensions, gregarisation and hunger, as well as the environmental variable of food. A full list of parameters and functions derived in this section can be found in Table 1 and Table 2, respectively. Finally, for simplicity we assume that the individual state and environmental components of *f*_*n*_ and *f*_*l*_ are separable, i.e.

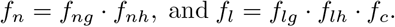

**Table 1:** Parameters used in numerical simulations.

| Variable | Description | Value | Units | Source |
| --- | --- | --- | --- | --- |
| $k$ | Normalised decay rate of serotonin | 0.9986 | hr<sup>-1</sup> | (26) [2]<br>[34] |
| $\kappa$ | Density of half normalised maximal serotonin release rate | 4.8293 | locusts·m<sup>-2</sup> | (28) [12] |
| $\delta$ | Normalised maximal serotonin release rate | 0.9960 | hr<sup>-1</sup> | (27)<br>[2][34] |
| $D$ | Linear diffusion coefficient | 0.01 | m<sup>2</sup>hr<sup>-1</sup> | |
| $\gamma$ | Non-linear diffusion coefficient | 4.5291 | locusts<sup>-2</sup> | (12) [13] |
| $A$ | Strength of non-local interactions | 6.83 | m<sup>3</sup>(hr·locust)<sup>-1</sup> | [45] |
| $r$ | Range of non-local interactions | 0.14 | m | [45][13] |
| $\nu$ | Normalised energy lost due to metabolism | 0.2496 | hr<sup>-1</sup> | [36] |

**Table 2:** Functions used in numerical simulations.

| Function | Description |
| --- | --- |
| $f_{ng} = A \left( 1 - \frac{2}{1 + e^{-15(n_g - 0.5)}} \right)$ | Function relating gregariousness to non-local interactions |
| $f_{lh} = 1.5 - n_h$ | Function relating hunger to non-local interactions |
| $f_c = e^{-c}$ | Function relating food density to local interactions |
| $\lambda(n) = 0.9362(2 - n)$ | The effect of locust feeding on satiation |
| $\psi(n) = 0.0384(2 - n)$ | The amount of food consumed based on locust hunger |

#### 4.1.1 The environment

In addition to locust densities, we include food resources in our model. Let *E* = *c*(***x***, *t*) denote the food density (mass of edible material per unit area). We assume that locust food consumption is based on their contact with food and their state, and on the time-scale of group formation food production is negligible, giving

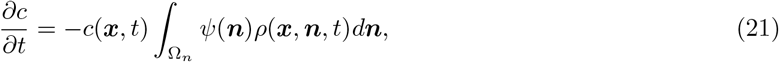

where *ψ*(***n***) is a function relating the locust’s state and food consumption rate. For the effect of food on movement we use the functional form from our previous study [22],

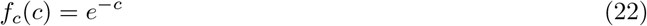

#### 4.1.2 Gregarisation

The most important state is that of gregarisation, we assume that locusts become increasingly gregarious as the local locust density increases (and solitarious with decreasing density). From Anstey et. al. [2], locust gregarisation is proportional to the amount of serotonin in their system, and this serotonin is released either through mechanosensory pathways (collisions) or cephalic pathways (sight and smell). We assume that this release is thus proportional to their local density but is bounded, and the serotonin decays at some metabolic rate. This results in equations given by

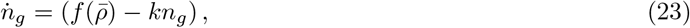

where, 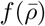 is a positive monotonically increasing bounded function representing normalised density dependant serotonin release rate, and *k* represents the normalised decay rate of serotonin in the locusts system.

For our function 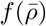 we select:

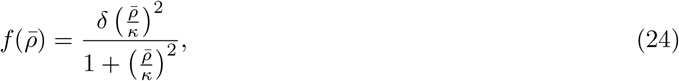

where *δ* is maximal normalised serotonin release rate and *κ* is the locust densities at which half this maximal rate occurs (this is the same function of gregarisation used by Topaz et. al. [45]). We note that we are only modelling the initial gregarisation process and this equation is not valid for locusts that have been gregarious for an extended period of time, i.e. its acute and not chronic gregarisation [29].

We then let *n*_*g*_ = 0.5 be the transition point between solitarious and gregarious and assume that this transition is fairly ‘quick’. We also assume that this only affects the non-local component of movement. Giving the gregarious component of *f*_*n*_ as a sigmoid function given by

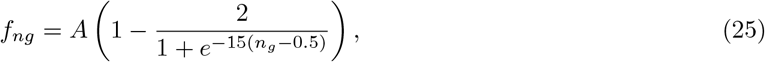

where *A* is the maximum strength of attraction/repulsion.

##### Estimating parameters

From (23), for a fixed 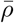 we can solve explicitly for *n*_*g*_(*t*) as

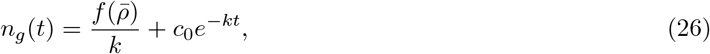

then we can find *k* by assuming *n*_*g*_(0) = 1, *n*_*g*_(*t*\*) = 0.05 and 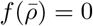, i.e. a gregarious locust fully solitarises (within 5% as our functions are asymptotic) after time, *t*\*. This gives

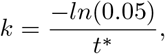

and letting *t*\* = 3 we get *k* = 0.9986. Next we can find *δ* by letting *n*_*g*_(0) = 0, *n*_*g*_(*t*\*) = 0.95 and 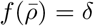. i.e. a solitarious locust will fully gregarise (within 5%) at the maximum rate of gregarisation after time, *t*\*. This gives

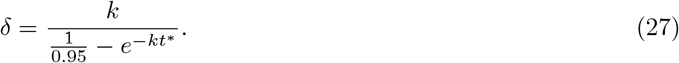

If again we let *t*\* = 3 (corresponding to symmetric gregarisation) we get *δ* = 0.9960. Finally, to find *κ*,we look at locust density prior to the onset of collective behaviour. We begin by setting 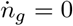, finding that

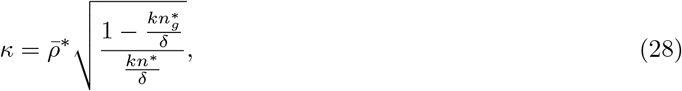

where 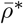 is the lower limit of collective behaviour (*≈* 5 from [12]) and 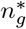 is state at which the stability condition from (15) is first satisfied. We can then find,

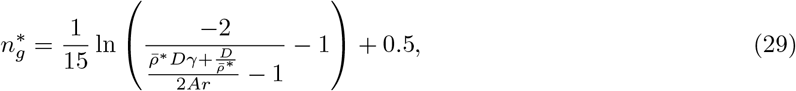

which gives 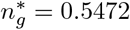, which in turn gives *κ* = 4.5351. We can then see gregariousness in time at different locust densities in Figure 3, compare with Figure 1C in [2] for gregariousness compared with treatment time.

**Figure 3.**
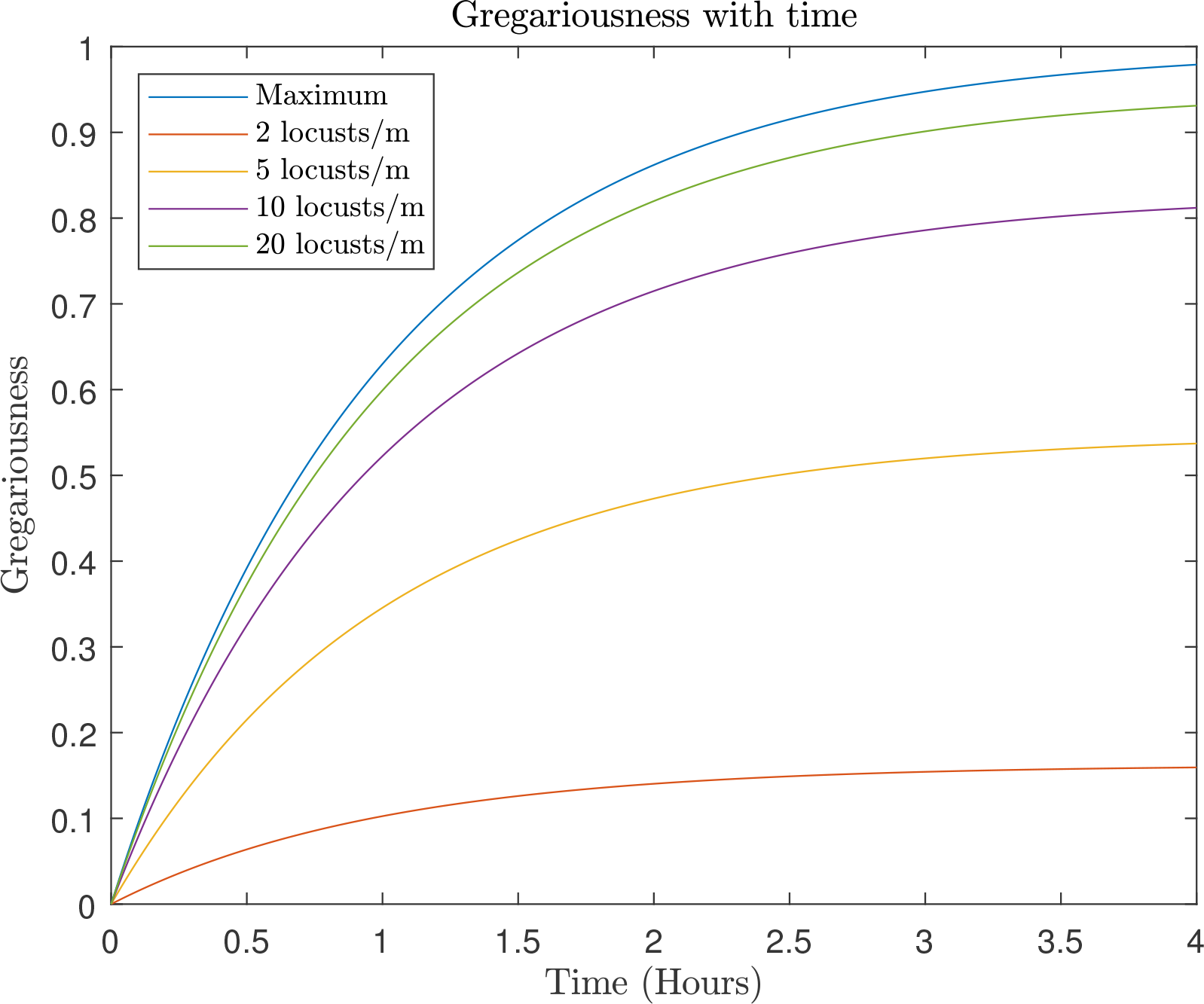
Gregarisation in time. Estimates of time required for a solitarious locust to reach various levels of gregarisation at different locust densities.

#### 4.1.3 Hunger

For the state dimension of hunger we assume that locusts become hungrier based on energy loss due to metabolism, and that locusts become satiated by eating at a rate proportional to their state. These assumptions give,

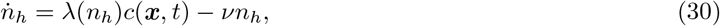

where *λ*(*n*_*h*_) describes how fast the locusts eat based on hunger, and *ν* is the energy lost due to metabolism. We then assume that hunger only affects local movement, with hungry locusts moving three times as fast as satiated ones [36] and the relationship is linear. Finally, we assume our previous parameter estimations correspond to a locust in hunger state *n*_*h*_ = 0.5. This gives,

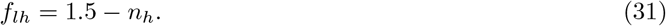

##### Estimating parameters

From (30) by assuming there is no food consumption, we get

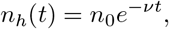

where *n*_0_ is our starting satiation. Then, if it takes some time, *t*\*, for the satiation level to go from 1 *→* 0.05, we can calculate *ν* as

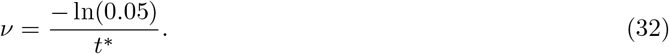

Based on [36], after 1 day of starvation locusts reached their maximum amount of locomotion if we let this be 12 hours of activity that results in *ν* = 0.2496. We then estimate *λ*(*n*), first by letting *λ*(*n*) be linear in *n*, i.e. *λ*(*n*) = *an* + *b* and our food be constant in time and space, we find

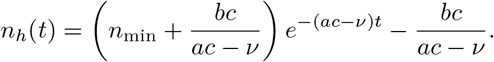

So if we assume a locust stays completely satiated after eating for 80 minutes in a 5 hour period [36] (i.e. *an* + *b* = 3.75*ν*) and a hungry locust eats twice as much as a satiated one we get

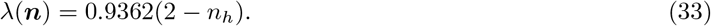

Finally, for *ψ*(***n***) in (21), we re-dimensionalise the parameter *κ* from our previous study [22], around *n*_*h*_ = 0.5 with the same linear relationship between hunger and consumption to get

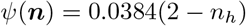

#### 4.1.4 Interpretation of analytic results in the context of locusts

Now that we have explicit forms of *f*_*n*_ and *f*_*l*_ we can interpret the analytic results derived in Section 3 as applied to locust foraging. From Section 3.1, we rewrite (11) using our states to find

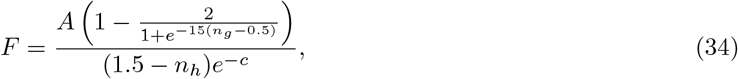

then from (9) and (13) we can see that hunger acts to decrease the maximum density of locusts by increasing their dispersal while the presence of food has the opposite effect (this relationship is inverted for the size of the support). Increasing gregarisation and attraction strength increases the maximum density of locust groups.

Using the results from Section 3.2. We can see from (70) and (76) that if all locusts are in the same state group formation occurs when

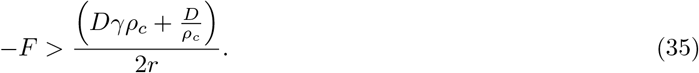

That is to say hungry locusts are required to be in a more gregarious state for group formation to occur. In addition food decreases the required gregariousness for group formation. These conditions carry over if the locusts can be considered as occupying two discrete states, with the fraction in the more gregarious state as the gregarious mass fraction. Then, using (20) we can see that: as the amount of available food increases the gregarious fraction required for group formation decreases. Additionally, the numerator in (20) is only in terms of ***n***_*s*_, thus as the solitarious locusts become hungrier (or more solitarious) this will increase the gregarious mass fraction required for group formation, with the presence of food acting to decrease it. Next, the denominator depends on the difference between states, thus if the gregarious state has less local movement than the solitarious state this decreases the gregarious mass fraction required for group formation and vice-versa. i.e. if all the gregarious locusts are satiated while the solitarious are not this would again decrease the mass fraction required. Finally, as 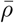 increases (or goes to 0) the gregarious fraction required for group formation increases suggesting an upper (and lower) locust density in order to form groups.

Of note, if only food is considered, this reduces to the stability condition found in our previous work [22], and if only the non-local forces are considered it reduces to that of Topaz et al. [45].

### 4.2 Numerical experiments

Using the parameters and functional forms defined above, Table 1 and Table 2 respectively, we perform numerical experiments to investigate the effect of states on group heterogeneity and the interaction between states and food distributions on group formation. We begin with simulations only including a gregarisation state, then including gregarisation and hunger. For all simulations, our spatial domain is the interval *x* = [0, *L*], where *L* = 3 (this is the same domain used by [45]), with periodic boundary conditions (i.e., *ρ*(0, ***n***, *t*) = *ρ*(*L*, ***n***, *t*)), for a total of 10 hours. These experiments are equivalent to those in our previous work [22], and the numerical scheme used is given in Appendix C.

The initial locusts density is given by

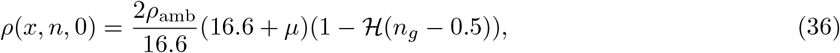

where *ρ*_amb_ is a ambient locust density, *µ* is some normally distributed noise, *µ* ∼ *𝒩* (0, 1), and *ℌ* is the Heaviside function, i.e. we have locusts uniformly distributed over the solitarious region of the gregarisation state. To ensure that simulations are comparable, we set-up three initial locust conditions and rescale them for each given ambient locust density, we then record the simulation with the highest final peak locust density. Finally, the initial condition for food is given by a smoothed step function of the form,

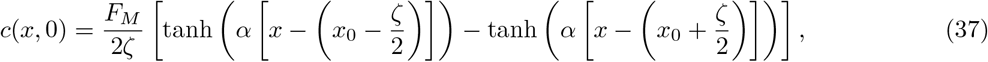

with *α* = 21, *x*_0_ = *L/*2, *F*_*M*_ being the food mass and *ζ* being the initial food footprint. We also introduce *ω* = 100*ζ/L* which expresses the size of the food footprint as a percentage of the domain.

#### 4.2.1 Gregarisation only

We can simulate just gregarisation with food, we will assume that the locusts have a hunger level of *n*_*h*_ = 0.5. Using (2), gives

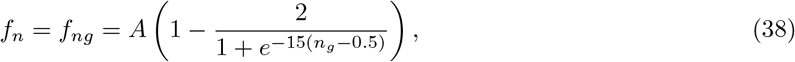

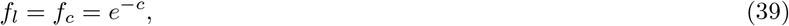

with (23) leading to

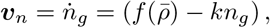

with 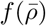 given by (24). Finally, food consumption, (21), is simplified to

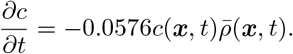

##### The effect of food on group formation

We begin by considering ambient locust densities from *ρ*_amb_ = 4.5 to *ρ*_amb_ = 5.2 and food footprint ranges from *ω* = 0% to *ω* = 50%. The results can be seen in Figure 4, with the highest peak density of locusts (max 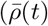) on the left and the final peak density (max 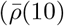) on the right. In contrast to our previous work, we find that by only considering gregarisation, food decreases the required density for group formation and that this effect is increased with decreasing food footprint and increasing food mass, as opposed to around some optimal width [22].

**Figure 4.**
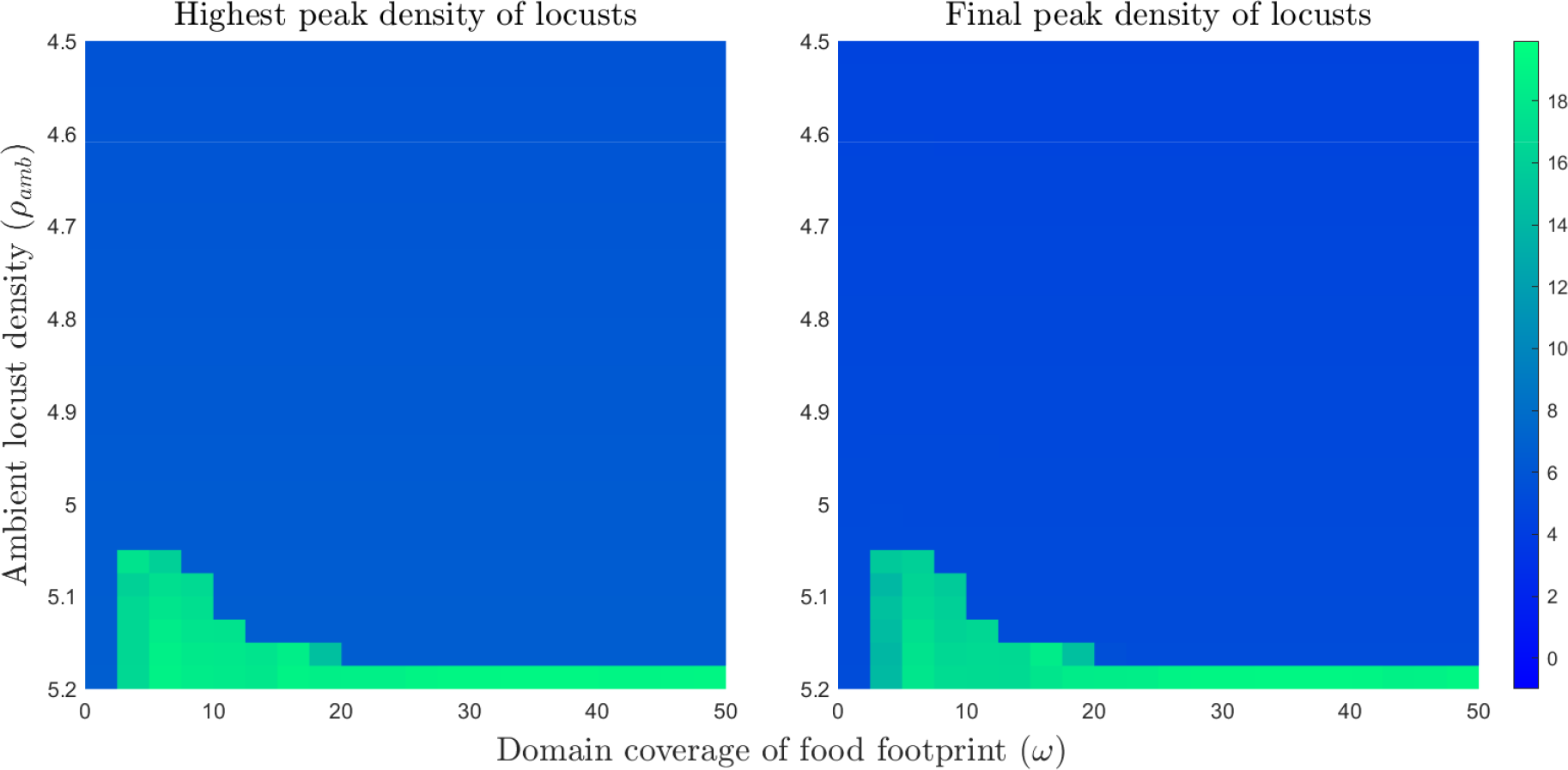
Maximum locust density with varying food footprint sizes and initial ambient locust densities. For the simulations, *x* = [0, 3] with periodic boundary conditions and *t* = [0, 12]. Ambient locust density ranges from *ρ*_amb_ = 4.8 to *ρ*_amb_ = 5.2, food footprint ranges from *ω* = 0% to *ω* = 50%. The plots show the maximum locust density for the varying food footprint sizes and ambient locust densities with a food mass of 1.

##### The effect of gregarisation on group heterogeneity

By looking at single simulations in Figure 5 we can see both the effect of gregarisation on group heterogeneity. In the plots, food is plotted in green with the total locust density given by a line that varies in colour based on the average locust gregarisation at that point, changing from blue being fully solitarious to red being fully gregarious. We can see that once a group has formed (as an aggregation with max density greater than initial ambient density) the most gregarious locusts end up in the middle of the group with a reduction in average gregarisation towards the edges.

**Figure 5.**
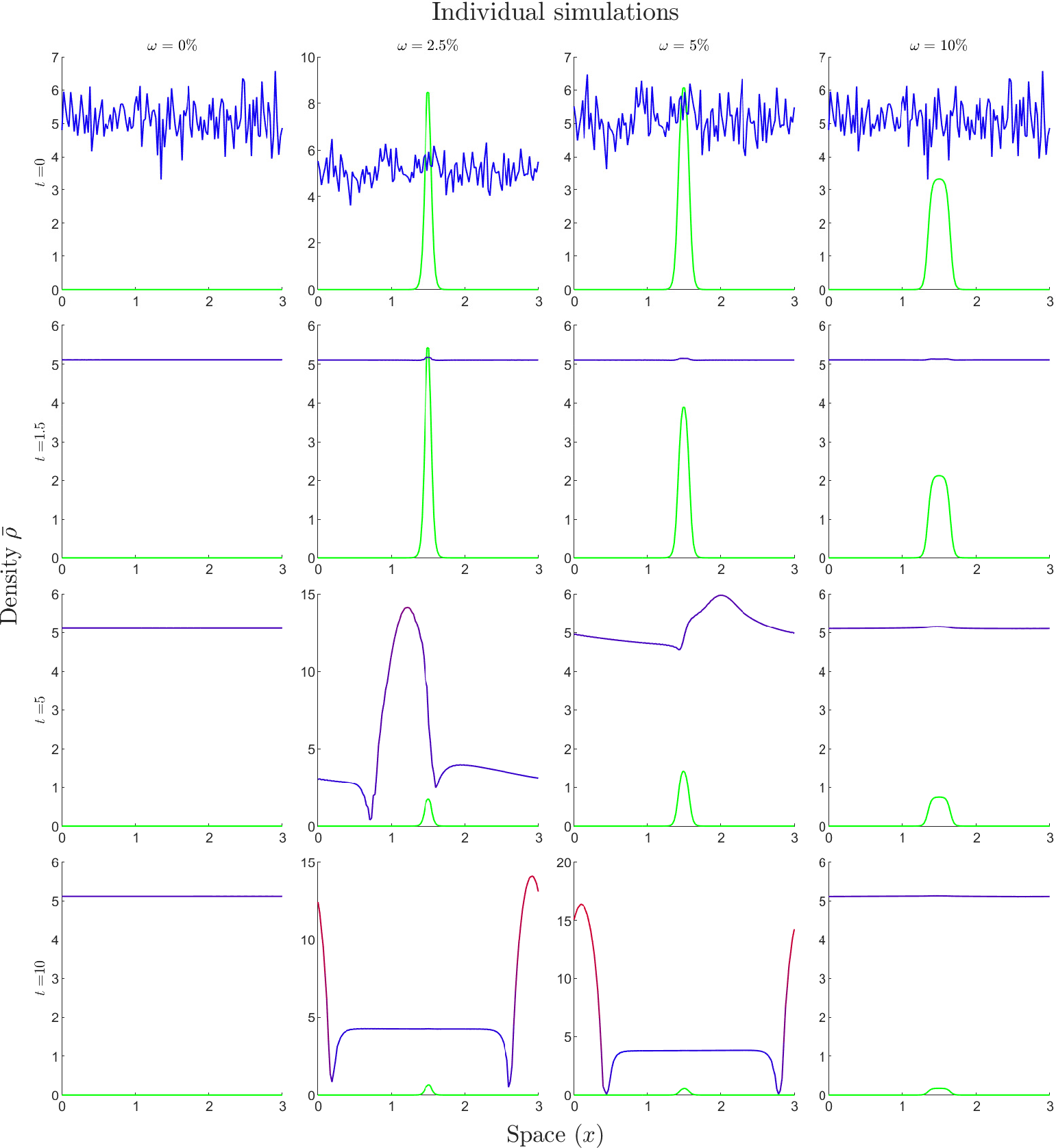
Individual simulations with varying food footprints. In the plots, food is plotted in green with the total locust density given by a line that varies in colour based on the average locust gregarisation at that point, changing from blue being fully solitarious to red being fully gregarious. For the simulations, ambient locust density is *ρ*_amb_ = 5.075, *ω* = 0%, 2.5%, 5%,10%, and *t* = 0, 1.5, 5, and 10.

#### 4.2.2 Gregarisation and hunger

We now repeat the previous experiment with the inclusion of hunger. Using (22), (25), and (31), the local and non-local forces become,

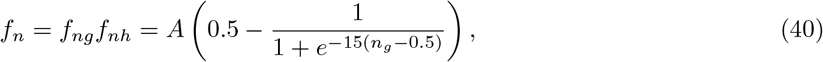

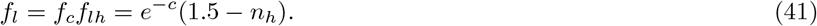

Next, the state space flux given by, (23) and (30),

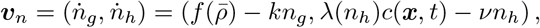

with 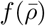 given by (24) and *λ*(*n*_*h*_) given by (33). Finally, food consumption, (21), is given by

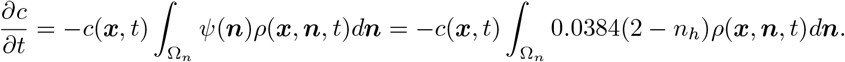

##### The effect of food on group formation with two internal states

We again consider ambient locust densities from *ρ*_amb_ = 4.5 to *ρ*_amb_ = 5.2 and food footprint ranges from *ω* = 0% to *ω* = 50%. The results can be seen in Figure 6, with the highest peak density of locusts (max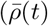 on the left and the final peak density (max(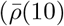 on the right. Most notably we see the re-emergence of the optimal food width, although less pronounced than in our previous work. In addition, the smallest food width of *ω* = 2.5% produces a high highest peak density of locusts at low ambient locust densities but these peaks are not maintained into the final peak density.

**Figure 6.**
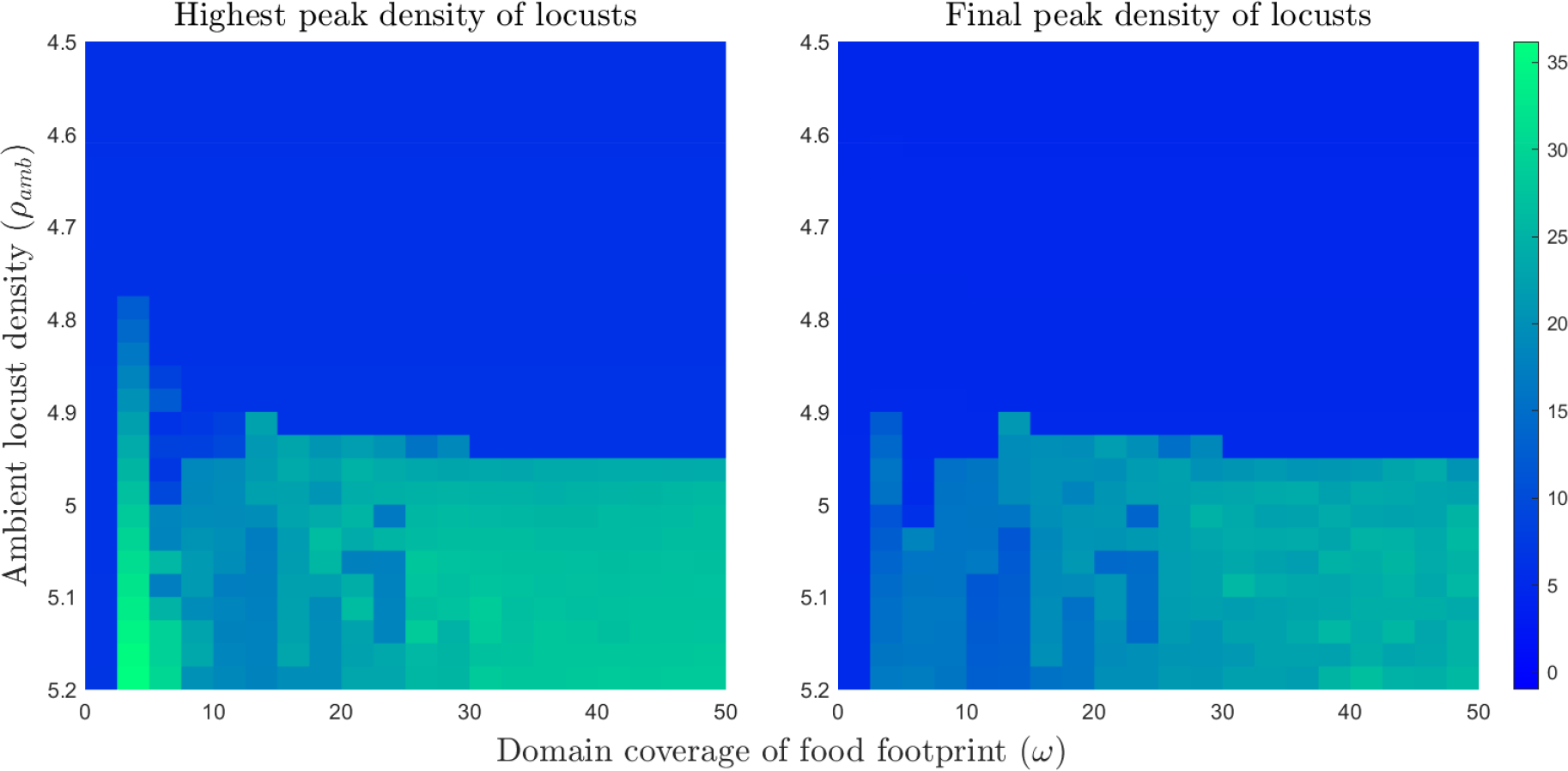
Maximum locust density with varying food footprint sizes and initial ambient locust densities. For the simulations, *x* = [0, 3] with periodic boundary conditions and *t* = [0, 10]. Ambient locust density ranges from *ρ*_amb_ = 4.8 to *ρ*_amb_ = 5.2, food footprint ranges from *ω* = 0% to *ω* = 50%. The plots show the maximum locust density for the varying food footprint sizes and ambient locust densities with a food mass of 1.

##### The effect of gregarisation and hunger on group heterogeneity

By looking at single simulations in Figure 7 we can see both the effect of gregarisation and hunger on group heterogeneity. In the Figure 7 food is plotted in green. Total locust density given by a line that varies in colour based on the average locust gregarisation at that point, changing from blue being fully solitarious to red being fully gregarious. Finally, the line width is the average hunger with thin being hungry and thick being satiated (however due to this potting technique there does appear an accordion style visual artefact). We can once again see the effect of the optimal food width. When the food is too narrow there is attempted group formation, however there are insufficient numbers of gregarious locusts so the group does not persist. If the food is too wide groups do not form. Finally, once a group has formed, on average the most gregarious and satiated locusts end up in the middle of the group with a reduction in gregarisation and an increase in hunger towards the edges. In addition, the most gregarious appear to be the most satiated, highlighting the existing relationship between gregarisation and foraging [21, 24].

**Figure 7.**
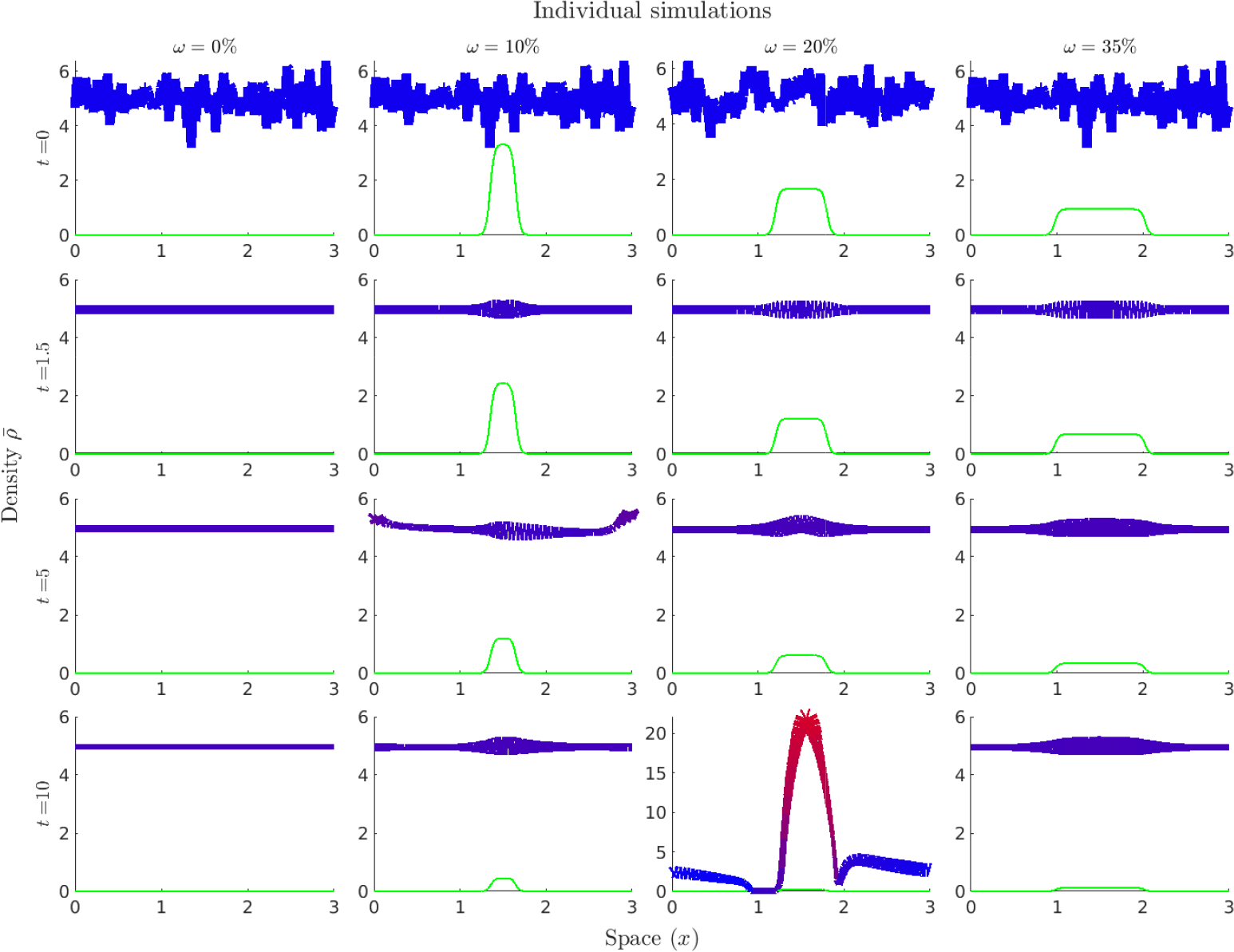
Individual simulations with varying food footprints. In the plots, food is plotted in green with the total locust density given by a line that varies in colour based on the average locust density at that point, changing from blue being fully solitarious to red being fully gregarious. For the simulations, ambient locust density is *ρ*_amb_ = 4.925, *ω* = 0%, 10%, 20%,35%, and *t* = 0, 1.5, 5, and 10.

We can next investigate the simulations with very narrow food sources, these results can be seen in Figure 8. In each simulation we see a high density of locusts in the centre of the food source, this is likely due both satiation and food increasing maximum locust density. From this state there are attempted group formations, at the lower densities these fail, but at the higher densities these are successful. However of note, in the final time-step the peak densities are about half that of the highest peak density, suggesting that hunger plays a role in the dispersion of locust groups.

**Figure 8.**
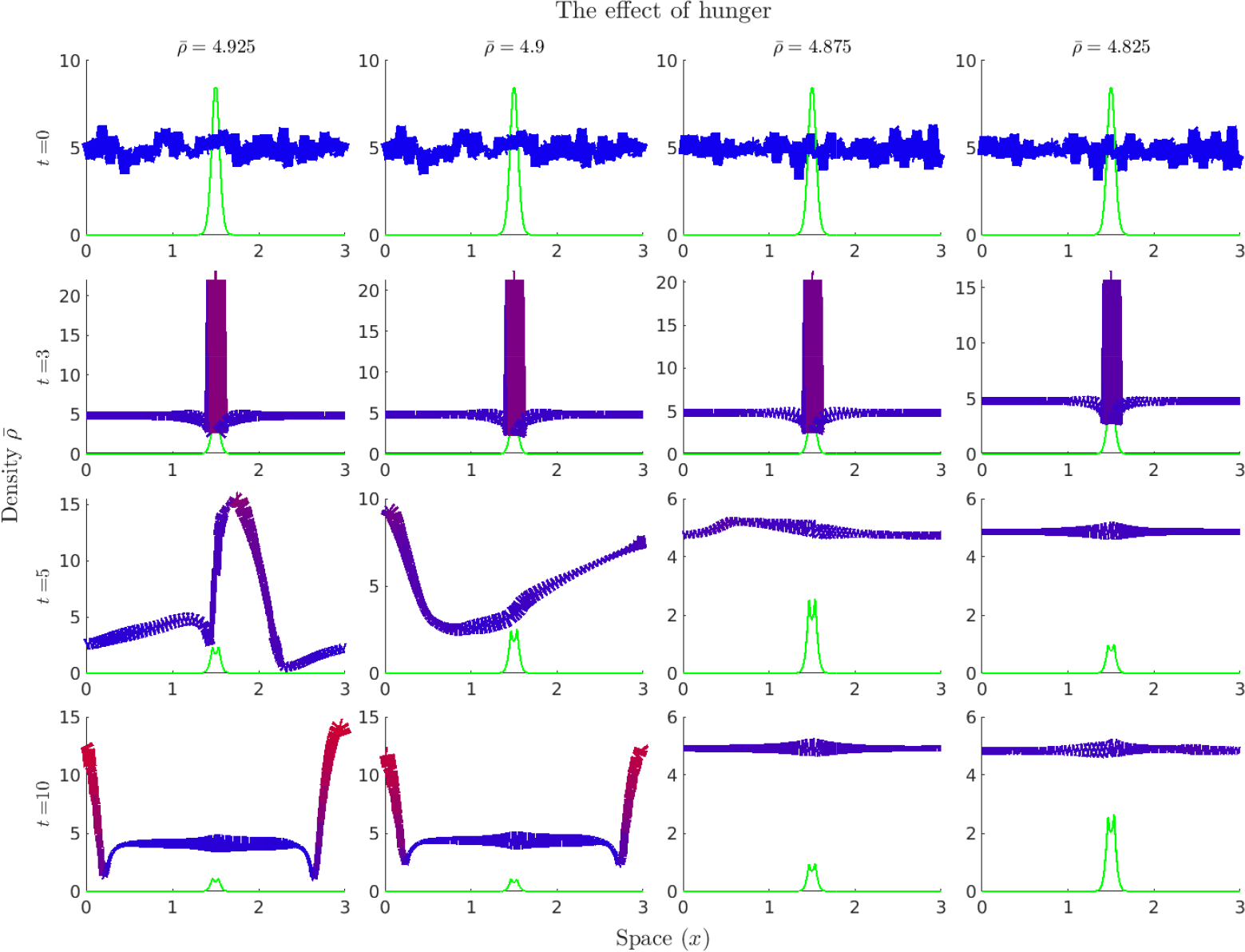
Individual simulations with varying ambient densities. In the plots, food is plotted in green with the total locust density given by a line that varies in colour based on the average locust density at that point, changing from blue being fully solitarious to red being fully gregarious. For the simulations, ambient locust density is *ρ*_amb_ = 4.925, 4.9, 4.875, 4.825, *ω* = 2.5%, and *t* = 0, 3, 5, and 10.

##### The effect of hunger on group dispersal

Finally, we can test the effect of hunger on group dispersal. In these experiments: we took a mass, *M* from 1 to 10, of locusts with an initial gregariousness of *n*_*g*_ = 0.95 both with and without a hunger state dimension (when hunger is included *n*_*h*_ = 0.95 initially). We placed them at the centre of the domain, and ran the simulation to *t* = 10. Finally, we measure the peak density of the locusts as well as the mean gregariousness at *t* = 10. The results can be seen in Figure 9, we found that hunger acts to both decrease the final peak density as well as the mean gregariousness leading to a faster breakdown in group cohesion. When *M ≤* 5 even in the absence of hunger the locust group dispersed, whereas with hunger we saw dispersal with *M ≤* 6.

**Figure 9.**
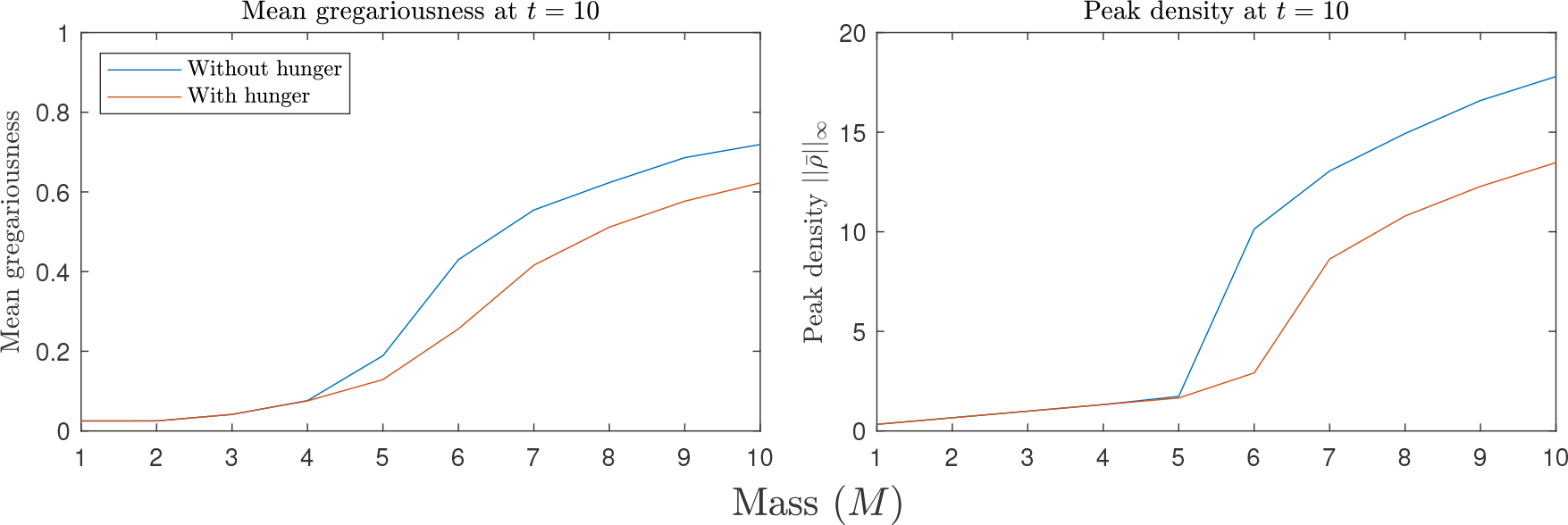
The effect of hunger on locust dispersal. In the left plot, is the average gregariousness with respect to mass and the right plot is the peak density (at time *t* = 10). In both plots, the red line includes hunger while the blue line does not. We can see that hunger has lead to a greater loss of gregariousness and a reduction in the peak density thus leading to an increase in locust dispersal.

## 5 Discussion

Heterogeneity plays an important role in group structure and dynamics [27]. This heterogeneity arises due to slow changing attributes such as age, size, personality, and/or fast-changing such as energy reserves and local environmental conditions. For example, hungrier individuals tend to migrate towards the front of moving groups [25, 30], with faster individuals holding a leadership position in flocks [35]. However, in the interest of mathematical tractability continuous models typically assume a mostly homogeneous approach to organism behaviour. In this paper, we have introduced a continuous kinematic model that takes into account both organism and environmental heterogeneity. Despite the increased complexity introduced by including heterogeneity we still gained some novel insights through analytic and numerical analyses.

Within the limitations of our assumptions in Section 3, we were able to derive several results. Firstly, we found that any change in state, ***n***, that increases the rate of local movement without a corresponding increase in non-local movement would decrease the maximum density of aggregations and thus increase the size of the support (and vice versa) at both large and small masses. In addition, any change in state that increases the rate of non-local movement compared to local movement would increase the maximum density of aggregations and thus decrease the size of the support. For these aggregations to form, if all organisms are in one state, then non-local (attractive) forces simply need to be greater than local forces for aggregations to form. We then considered two possible states, termed solitarious and gregarious in relation to the strength of the non-local attraction. In environmental conditions that reduce dispersal the gregarious fraction required for group formation decreases. In regards to states, states that increase the local and non-local terms will increase the gregarious mass fraction required for aggregation formation. Locally, if the gregarious state has less local movement than the solitarious state this decreases the gregarious mass fraction required for group formation and vice-versa. Non-locally, decreasing the non-local force of both the gregarious and solitarious states decreases the gregarious mass fraction required for group formation and vice-versa. Finally, as 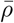 increases the gregarious fraction required for aggregation formation increases suggesting an upper organism density limit for this process.

We then applied the model to locust foraging including food interaction, and state dimensions of hunger and gregarisation. Within this context, in Section 4.1.4 we interpreted our analytic results. We firstly found that hunger acts to decrease the maximum density of locusts by increasing their dispersal while the presence of food has the opposite effect (this relationship is inverted for the size of the support). Secondly, increasing gregarisation and attraction strength increases the maximum density of locust groups. In addition, food decreases the required gregariousness for group formation. These conditions carry over if the locusts can be considered as occupying two discrete states, with the fraction in the more gregarious state as the gregarious mass fraction. We saw that: as the amount of available food increases the gregarious fraction required for group formation decreases. Additionally, as the solitarious locusts become hungrier (or more solitarious) this increases the gregarious mass fraction required for group formation, with the presence of food acting to decrease it. Next, if the gregarious state has less local movement than the solitarious state this decreases the gregarious mass fraction required for group formation and vice-versa. i.e. if all the gregarious locusts are satiated while the solitarious are not this would again decrease the mass fraction required. Finally, as 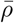 increases (or goes to 0) the gregarious fraction required for group formation increases suggesting an upper (and lower) locust density in order to form groups.

Next, our numerical results from Section 4.2 showed that considering states has a variety of effects on group formations and heterogeneity. By considering only a continuous representation of gregarisation, we found that once a group has formed the most gregarious locusts end up in the middle of the group with a reduction in average gregarisation towards the edges. Of note, the previously found optimal food width was no longer present, instead as food became more concentrated the density of locusts required for group formation decreased. However, once we included the state dimension of hunger, we once again saw the optimal food width effect. When the food is too narrow there is attempted group formation, however there is an insufficient mass of gregarious locusts so the group does not persist. Alternatively, if the food is too wide no attempt at group formation occurs.

It has been observed in many different species that individuals position themselves based on nutritional requirements, with hungrier individuals migrating towards the edges of the group and increasing the average distance to their neighbours [25, 30, 37]. We saw that once a group has formed, on average the most gregarious and satiated locusts end up in the middle of the group with a reduction in gregarisation and an increase in hunger towards the edges. Our results with modelling hunger suggest a rather simple mechanism for this observed behaviour. In that, hunger increasing local dispersive movement drives the hungry individual to the edges of the group and has the additional effect of lowering the local maximum density. In addition, the most gregarious appear to be the most satiated, highlighting the advantage gregarisation offers when foraging [20, 21, 22, 24].

In addition, we observed that hunger may lead to a breakdown in group cohesion. By increasing the dispersion, this leads to a lowering of maximum density, which in turn leads to a decrease in gregariousness. When this occurred with a sufficiently small mass of locusts it lead directly to group dispersal. We do note that we haven’t included hunger dependant cannibalism which may operate to counteract this effect.

Even with the more complex state model, we did require some simplifying assumptions that limit the direct biological relevance of the model. In addition, in order to perform the numerical experiments we were limited in the number of simultaneously simulated state dimensions due to computational complexity. Further explorations within the context of locust foraging could look at changing our state assumption, that the effect of internal states and the environment were separable and relatively simple, or adding additional states and interactions. Some examples for extension are a more complex hunger and feeding interactions [41], including cannibalism [5], or the inclusion of a heterogeneous age structure. Finally, while we have focused mainly on locusts and foraging, the modelling technique itself could potentially be used to explore a variety of different scales from microscopic to macroscopic [23, 31].

## A Derivation of local-flux

Following the work of Painter and Sherratt [33], we derive the local interactions as the limit of a lattice model,

We begin here by considering organism movement on a (*N* + 1)-dimensional lattice with one dimension of space and *N* dimensions corresponding to the organisms internal state with random movement restricted to the spatial dimension and the state space traversed by some advective velocity. Let each dimension, *d*, of the internal state have *m*_*d*_ possible choices, with an arbitrary state at the indexed location, ***j***, given by the *N* dimensional vector ***n***_***j***_. Let 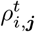 be the number of organisms at site *i* and with internal state ***n***_***j***_, at time *t*, and 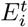 be the environmental conditions at site *i*. In addition let,

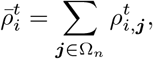

i.e. 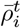 is the total number of organisms at spatial site *i* and time *t* regardless of state.

We assume that the transition probabilities for a organism at the (*i*, ***j***)^*th*^ site depends on the environmental conditions at that site, the state of the organism, and the relative population density between the current site and neighbouring sites. If we let 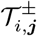 be the probability at which organisms at site (*i*, ***j***) move to the right, +, and left, *−*, during a timestep, then our transition probabilities are

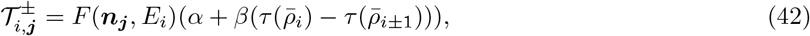

where *F* is a function of environmental conditions and state, *τ* is a function related to the local organism density, and *α* and *β* are constants. Finally, individuals transition around the state-space given by the boundary fluxes

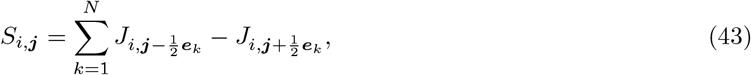

where ***e***_*k*_ is the unit state-space vector in the arbitrary direction *k*. We also assume that individuals either move in space or change internal state at each time step. Then the number of individuals at site (*i*, ***j***) at time *t* + Δ*t* is given by

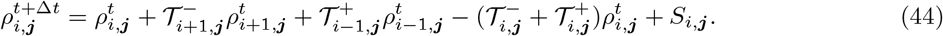

Substituting (42) and (43) into (44) gives

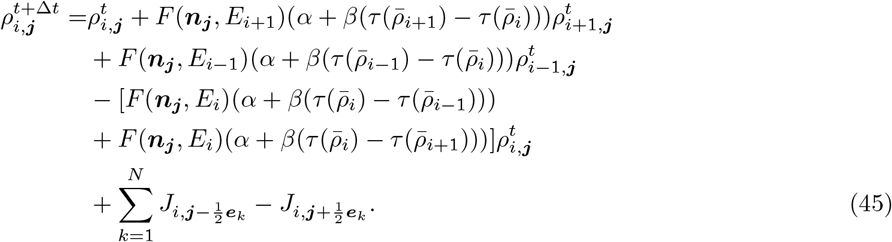

We the rearrange (45) to take out the common factors *α* and *β*, giving

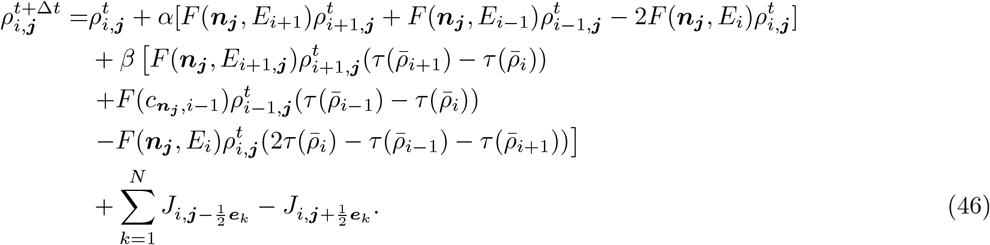

We then Taylor expand the terms in (46) to obtain the equation in relation to the site (*i*, ***j***) at time *t* only. Beginning with,

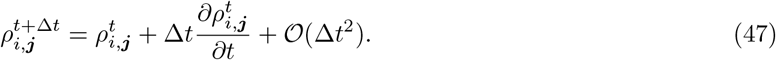

Then for the terms related to *α* we get

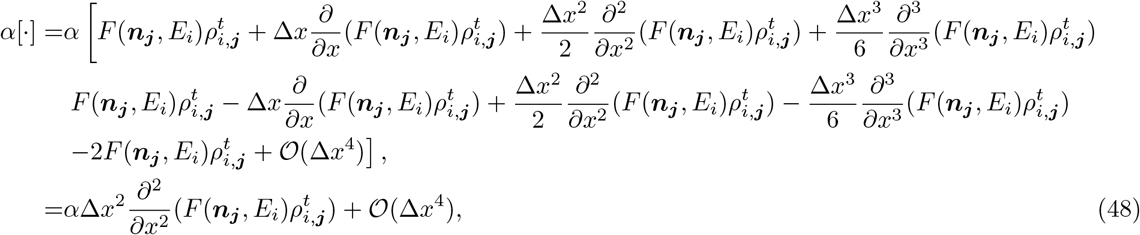

as the 0^*th*^, 1^*st*^, and, 3^*rd*^ order terms of Δ*x* cancel each other out. We then turn our attention to our terms involving *β*, we will Taylor expand each multiplication individually as otherwise the terms become unimaginably unmanageable. To begin,

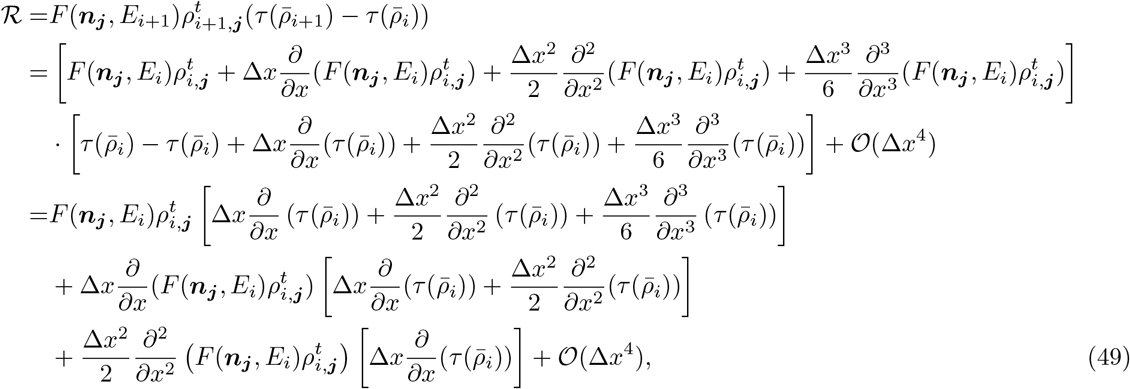

and

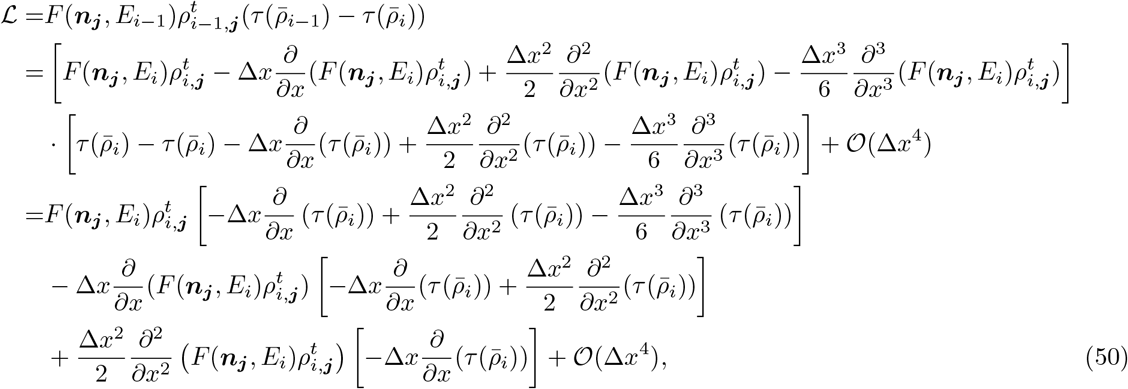

and finally,

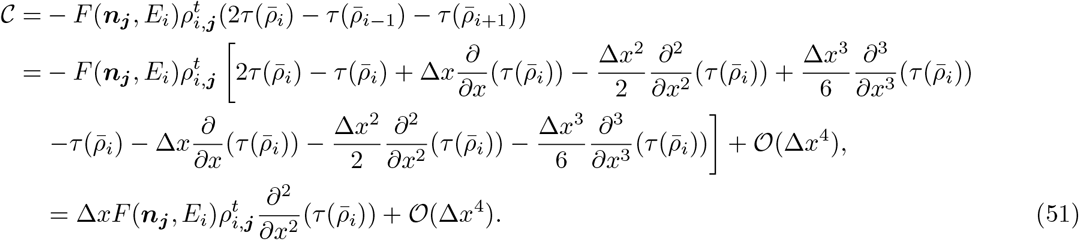

Adding (49), (50), and (51), gives

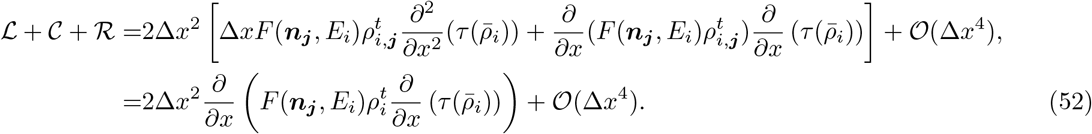

Lastly, for our flux around the state-space we get

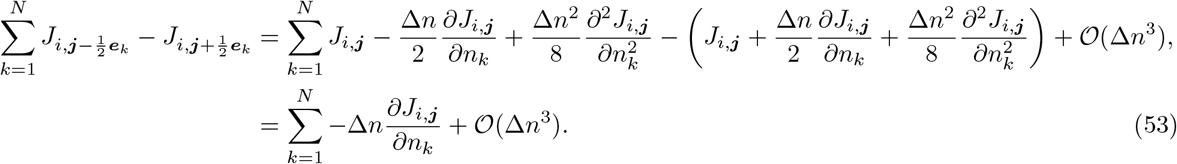

Combining (47), (48), (52), and (53) into (46), gives,

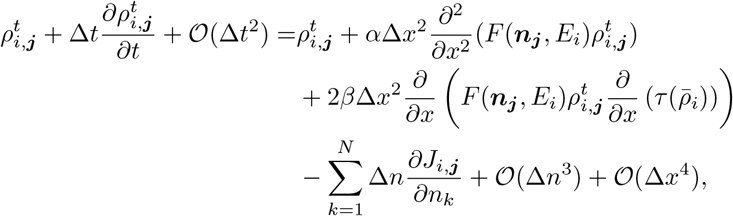

which we rearranging to obtain

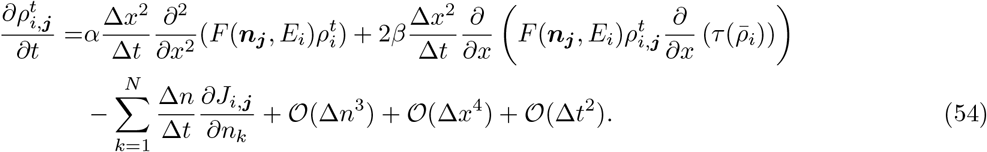

We then substitute our functions,

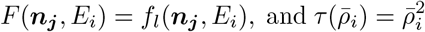

to obtain

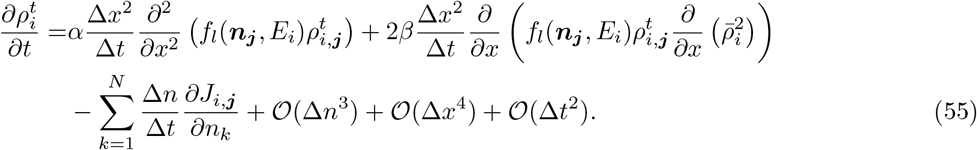

We then take the limit as Δ*x*, Δ*t*, Δ*n →* 0 such that,

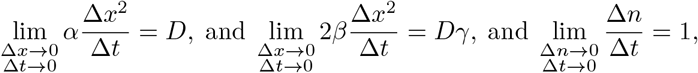

to find,

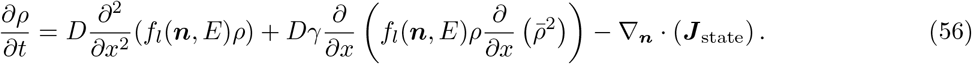

Which we then rearrange to find our local flux as

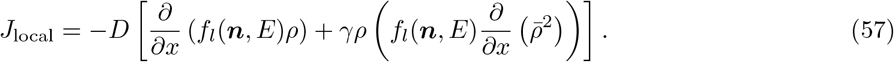

## B Full analytic results

In this appendix we present the calculations for our analytic results.

### B.1 Density of aggregations

In order to facilitate analysis, we begin by introducing some simplifying assumptions. Under which we can estimate the maximum density and width of a aggregations at both the large and small mass limits in one dimension. To begin, our assumptions are *E* is constant in space and time, and all the organisms are in the same unchanging state i.e. 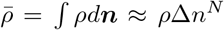 where *N* is the number of state dimensions and Δ*n* represents a small area in state space. Finally, we will label the support of *ρ* as Ω^*′*^. These assumptions allow is to rewrite (6) as,

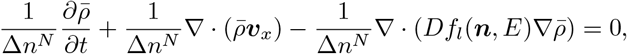

with

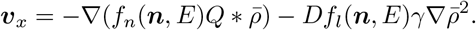

This can be rewritten as

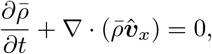

with

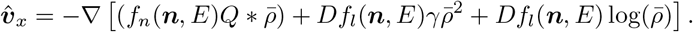

This is a gradient flow of the form

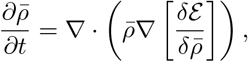

where

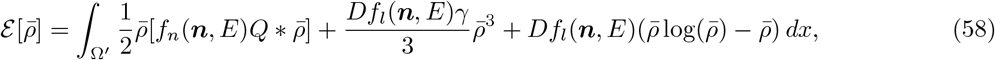

with the minimisers satisfying

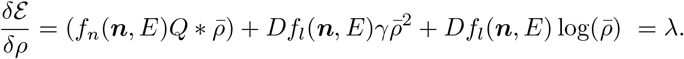

In the next sections, we follow the work of [8, 14, 43] and with further simplifying assumptions we consider the maximum density and support of aggregations with small and large numbers of organisms (i.e. as *M →* 0 and *M → ∞*, respectively). We term these the small mass limit and large mass limit, respectively.

#### B.1.1 Large mass limit

We begin with (58) and then further assume the following: 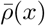 is approximately rectangular. Additionally, while the support of 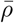, Ω^*′*^, is infinite due to the linear diffusion the bulk of the mass is contained as a series of aggregations, we will approximate the support of an aggregation as Ω. Finally, for a single aggregation we assume that the support is far larger than the range of *Q*. We thus approximate *Q* ≈ *VQδ*(*x*), where *δ*(*x*) is the Dirac delta function. We note that the *V*_*Q*_*δ*(*x*) needs to preserve the volume of *Q* so *V*_*Q*_ = *∫ Qd****x***. Then as 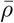 is rectangular

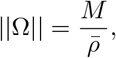

where *M* is given by (1). Substituting into (58) we get

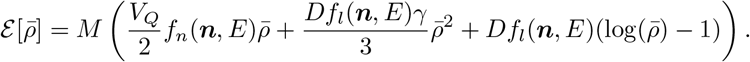

We can then find

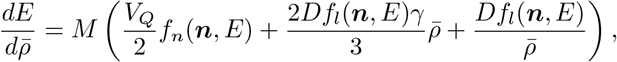

which has critical point at

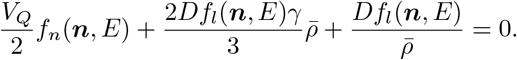

Thus

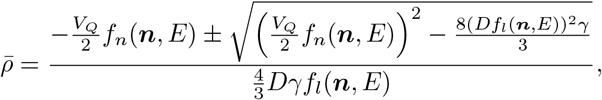

which simplifies to

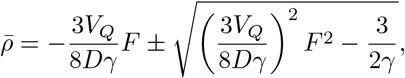

where

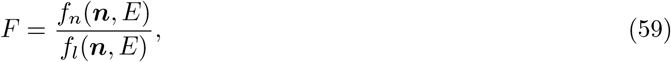

is the ratio of our non-local and local forces. We then take only the positive root, giving

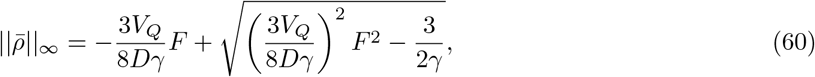

with support

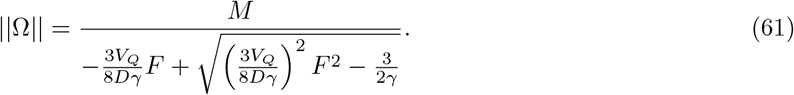

There are a few things to note in these equations. Firstly for aggregations to exist, i.e. 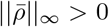, *F* must be less than 0 corresponding to an attractive social potential (or an attractive local movement; however, if both local and non-local components are attractive our original assumption of 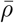 being rectangular would not hold as the equation is singular). Next, from this we find that any change in state, ***n***, that increases the rate of local movement without a corresponding increase in non-local movement would decrease the maximum density of aggregations and thus increase the size of the support (and vice versa). Finally, any change in state that increases the rate of non-local movement compared to local movement would increase the maximum density of aggregations and thus decrease the size of the support.

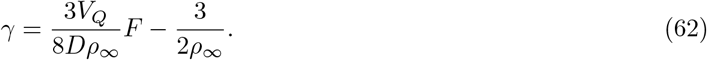

#### B.1.2 Small mass limit

Assuming that for a single aggregation we can approximate the social interaction potential using a Taylor expansion. In this section we use *Q*(*x*) given by (7), with 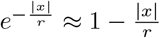. Additionally, we ignore the effect of linear diffusion within Ω, giving (58) as

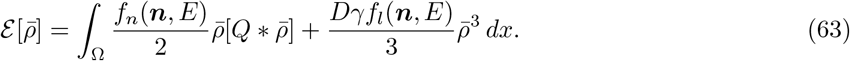

Based on these assumptions we can find

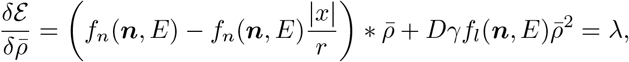

which becomes

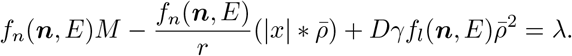

We then exploit the property that (|*x*|)_*xx*_ = 2*δ*(*x*) and differentiate twice to obtain

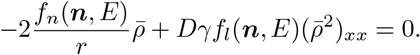

Following [8] we place the maximum of 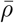 at the origin; this implies 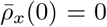 and 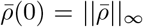. We then let,

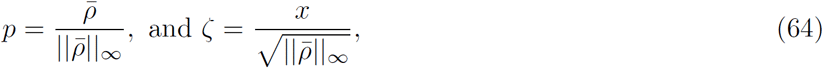

giving,

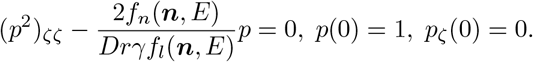

We then multiply through by (*p*^2^)_*ζ*_ and integrate to obtain,

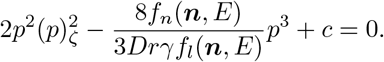

Then applying the conditions at *ζ* = 0 we find,

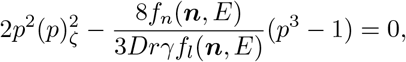

which can be simplified to,

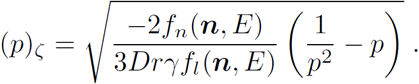

Performing a separation of variables gives

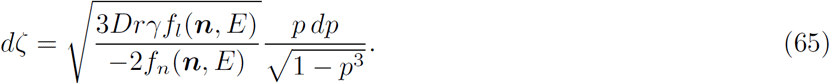

We can then find the implicit solution,

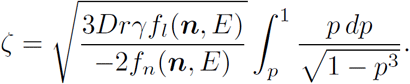

As *p* →0, 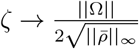, giving

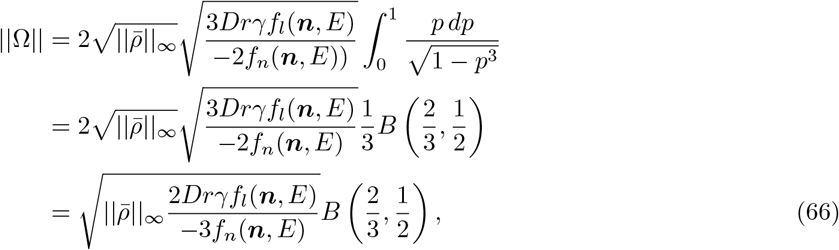

where *B* is the *β*-function (for definition see [50], page 207). Next using the mass constraint,

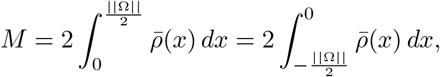

and substituting (64) we obtain

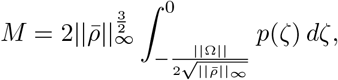

which using (65) becomes,

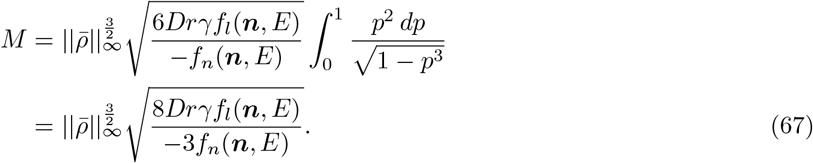

Then using (66) and (67) we can find ∥Ω∥ and ∥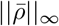∥_*∞*_ in terms of *M* and *F* ((1) *and (11), respectively)* giving

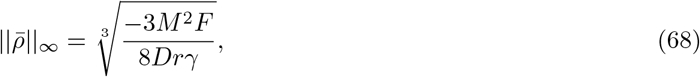

and

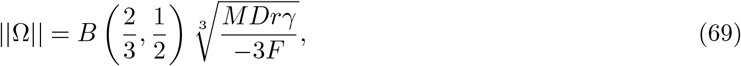

which gives a similar relationship as the large mass limit.

#### B.1.3 Comparison of mass limits and simulations

We can check the accuracy of our estimates by comparing them to simulation results (Details of the numerical scheme can be found in Appendix C). We begin by defining two dimensional state space, *(n*_1_, *n*_2_), with the first dimension only affecting the non-local component of force and the second dimension affecting the local component. For example, these could be considered in locusts to be dimensions of gregarisation and hunger, respectively. We first let,

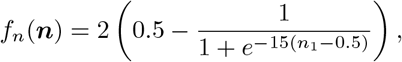

and

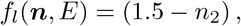

in addition we let,

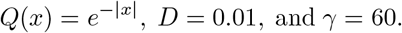

For each simulation we place a mass of organisms, *M*, in every combination of the states *n*_1_ = 0.75, 0.85, 0.95 and *n*_2_ = 0.15,0.55,0.95 in the center of the domain (we vary the domain size according to mass so there is no boundary interaction) and run it to a pseudo steady state, *t* = 1000.

The results for 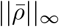 can be seen in Figure 10 and the results for ∥Ω∥ can be seen in Figure 11. In the plots, the dotted lines represent the minimum estimates between the small and large mass limits, and the solid lines represent simulated results. As ∥Ω∥ is theoretically infinite due to the linear diffusion, for the simulated ∥Ω∥ we select the region for which 98% of the mass, *M*, is contained. We can see that as *M* increases the simulated limits approach those given by the theoretical estimates, however there is some error likely due to error in the simulations and our method of approximating ∥Ω∥. In addition, the small mass estimates are considerably less accurate than the large mass limits (due to ignoring linear diffusion), however, the estimates display qualitatively similar behaviour.

**Figure 10.**
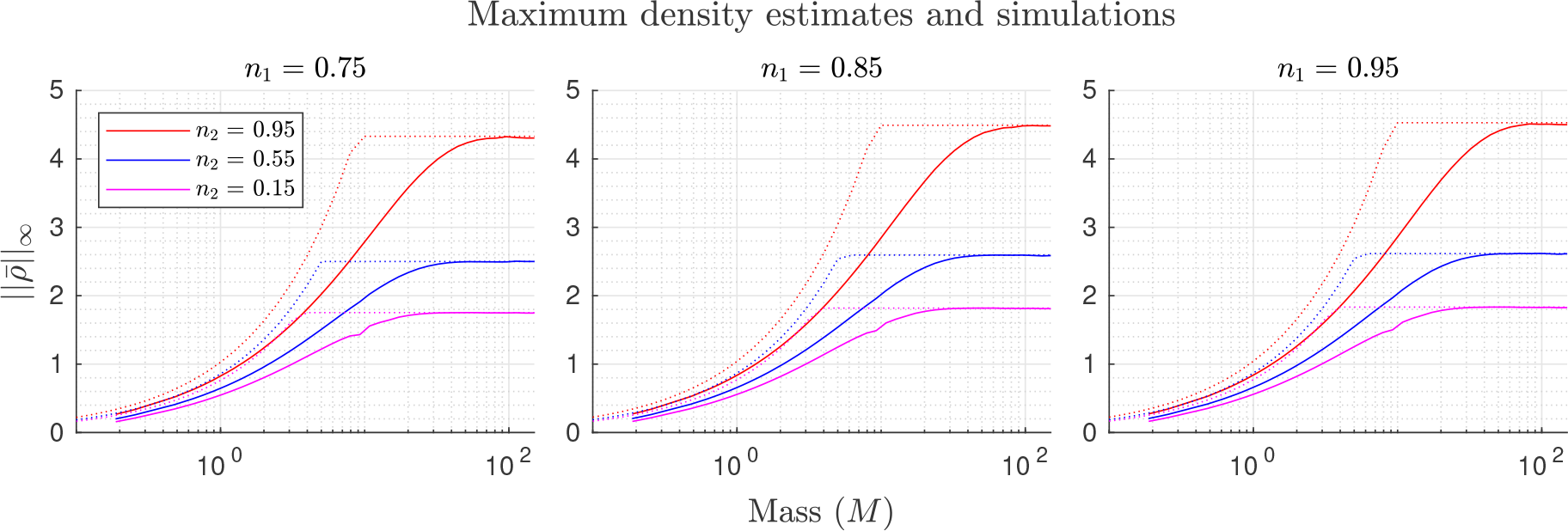
Small and large mass density estimates and simulations. Estimates for the small and large mass density limits in different states are plotted in dotted lines, with the corresponding simulation results plotted in solid lines.

**Figure 11.**
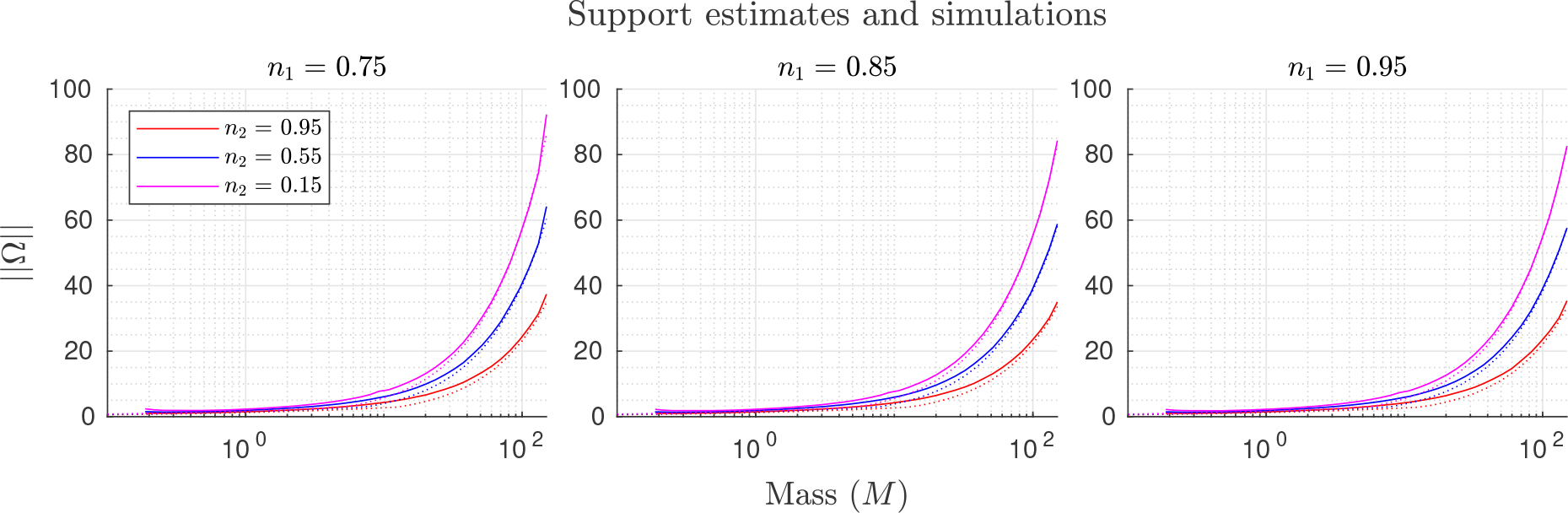
Small and large mass max density and support, estimates and simulations. Estimates and simulations for the maximum density (left) ans size of the support (right). Estimates for different states are plotted in dotted lines, with the corresponding simulation results plotted in solid lines. As ∥Ω∥ is theoretically infinite due to the linear diffusion, for the simulated ∥Ω∥ we select the region for which 98% of the mass, *M*, is contained.

### B.2 Linear stability analysis of homogeneous steady states

In order to gain insights into the conditions under which aggregations can form, we investigate the stability of spatially-homogeneous steady states. In this analysis we perturb the homogeneous steady state by adding a small amount of noise. We then find under what conditions the small perturbations grow and are likely to lead to aggregations. We again assume that *E* is constant in space and time.

#### B.2.1 A single state

We begin by considering all organisms in a fixed single state, ***n***_*f*_, i.e. 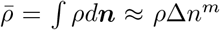 number of state dimensions. These assumptions give,

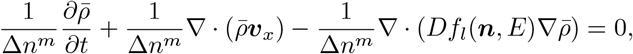

With

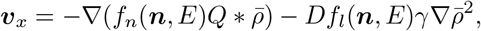

which then becomes

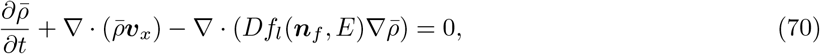

with

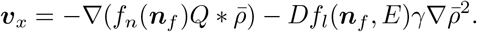

We then perturb around a homogeneous steady state, by letting 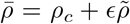 where 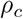 is the homogeneous steady state and 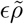 is a small perturbation (*ϵ≪*1). Substituting this into (70) and removing terms of *𝒪* (*ϵ*^2^), we find

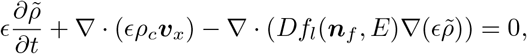

with

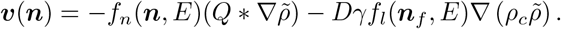

Next we take Fourier transforms in space and Laplace transforms in time of 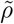, i.e. 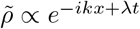 to obtain

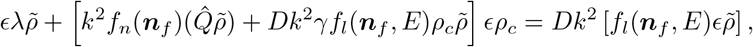

where 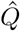 is the Fourier transform of *Q*. Then, dividing through by 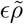 we get

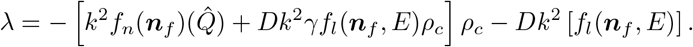

If *λ >* 0 then small perturbations will grow in time, we can then find the condition for instability in terms of our two state based force, as

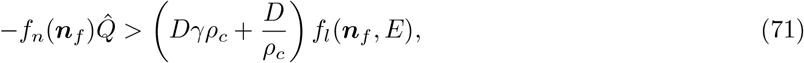

or alternatively

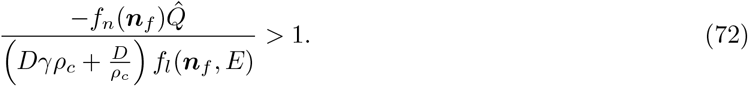

#### B.2.2 Two states

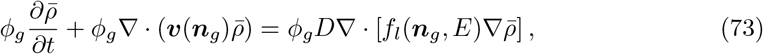

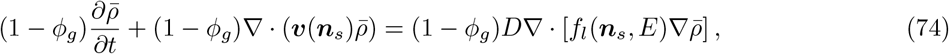

with

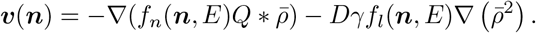

As in the single state case, we then perturb around a homogeneous steady state, by letting 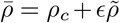 where *ρ*_*c*_ is the homogeneous steady state and 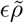 is a small perturbation (*ϵ ≪* 1). Substituting this into (73) and (74), removing terms of *𝒪* (*ϵ*^2^), then adding (73) and (74) together we find

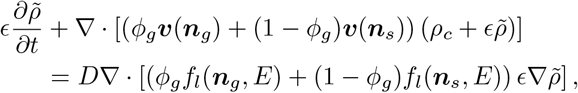

with

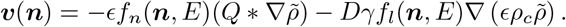

This can be simplified further into (by removing the next bunch of *𝒪* (*ϵ*^2^)),

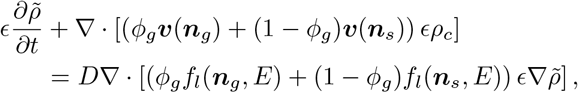

with

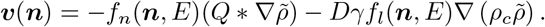

Next we take Fourier transforms in space and Laplace transforms in time of 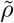, i.e. 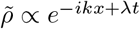 to obtain

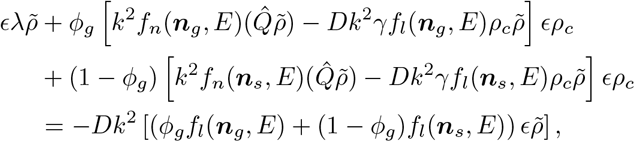

where 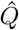 is the Fourier transform of *Q*. Then, dividing through by 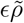 we get

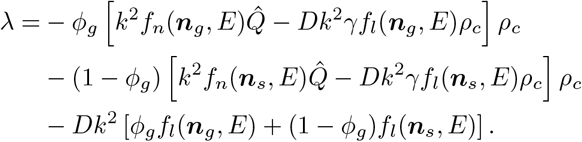

If *λ >* 0 then small perturbations will grow in time, we can then find the condition for instability in terms of *ϕ*_*g*_ as

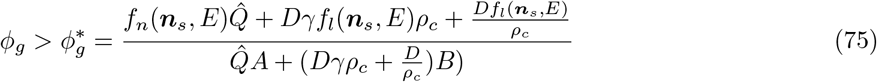

where

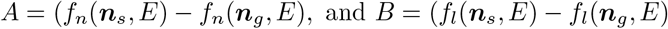

From this we can see that: in environmental conditions that reduce dispersal the gregarious fraction required for aggregation formation decreases. The numerator is only in terms of ***n***_*s*_, thus states that increase the local and non-local terms will increase the gregarious mass fraction required for aggregation formation. The denominator depends on the difference between the states: Locally, if the gregarious state has less local movement than the solitarious state this decreases the gregarious mass fraction required for aggregation formation and vice-versa. In addition, if the two local forces are equal, *f*_*l*_(***n***_*g*_) = *f*_*l*_(***n***_*s*_), this reduces to the stability condition found in our previous two population model [22], and if the local forces are 0 it reduces to that of Topaz et al. [45]. Non-locally, decreasing the non-local force of both the gregarious and solitarious states decreases the gregarious mass fraction required for aggregation formation and vice-versa (as *f*_*n*_ *<* 0 is attraction and *f*_*n*_(***n***_*g*_, *E*) *< f*_*n*_(***n***_*s*_, *E*)). Finally, as 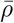 increases the gregarious fraction required for aggregation formation increases suggesting an upper organism density in order to transition away from the homogeneous steady state.

For our specific function 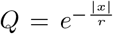, we begin by taking the one dimensional Fourier transforms of *Q* using the following definition,

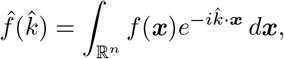

to get

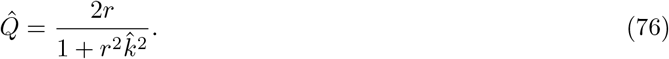

As 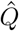 has a maximum value at 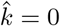, we let 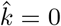 and substitute into (19), which gives,

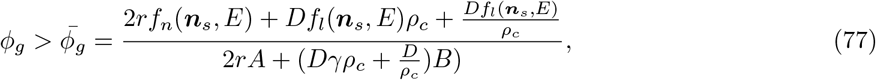

where

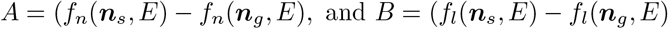

From this we can find the maximum homogeneous density, *ρ*_*c*_, that aggregations can still form. So taking (20) and substituting *ϕ*_*g*_ = 1 gives,

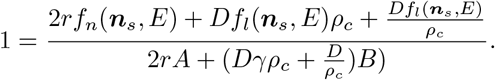

This gives

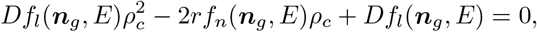

and this has solutions

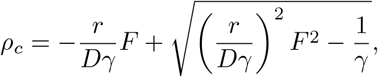

where *F* is given by (11) for the state ***n***_*g*_. Rather neatly, if 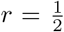, corresponding to 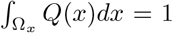, this becomes

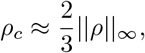

where ∥*ρ*∥_*∞*_is given by (9), this is similar to the relationship previously derived by us [22]. Finally, we note that the upper limit of aggregation formation only depends on the forces of organisms in the gregarious state.

## C Numerical Scheme

We now derive the numerical scheme for (6) in one spatial dimension using a finite volume method (FVM). In this section we use index notation inspired by Karlsen and Risebro [28], and Bürger et al. [15]. In this notation the arbitrary cell *i* is given by the tuple of vectors *i* = (*i*_***x***_, *i*_***n***_) with a spatial step indexed by *i ± e*_*x*_ and a state step indexed by *i ± e*_*n*_. This gives cell boundaries as 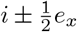 for space and 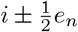 for state. We will also define

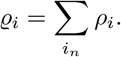

For the numerical scheme the terms are described in Table 3.

**Table 3:**
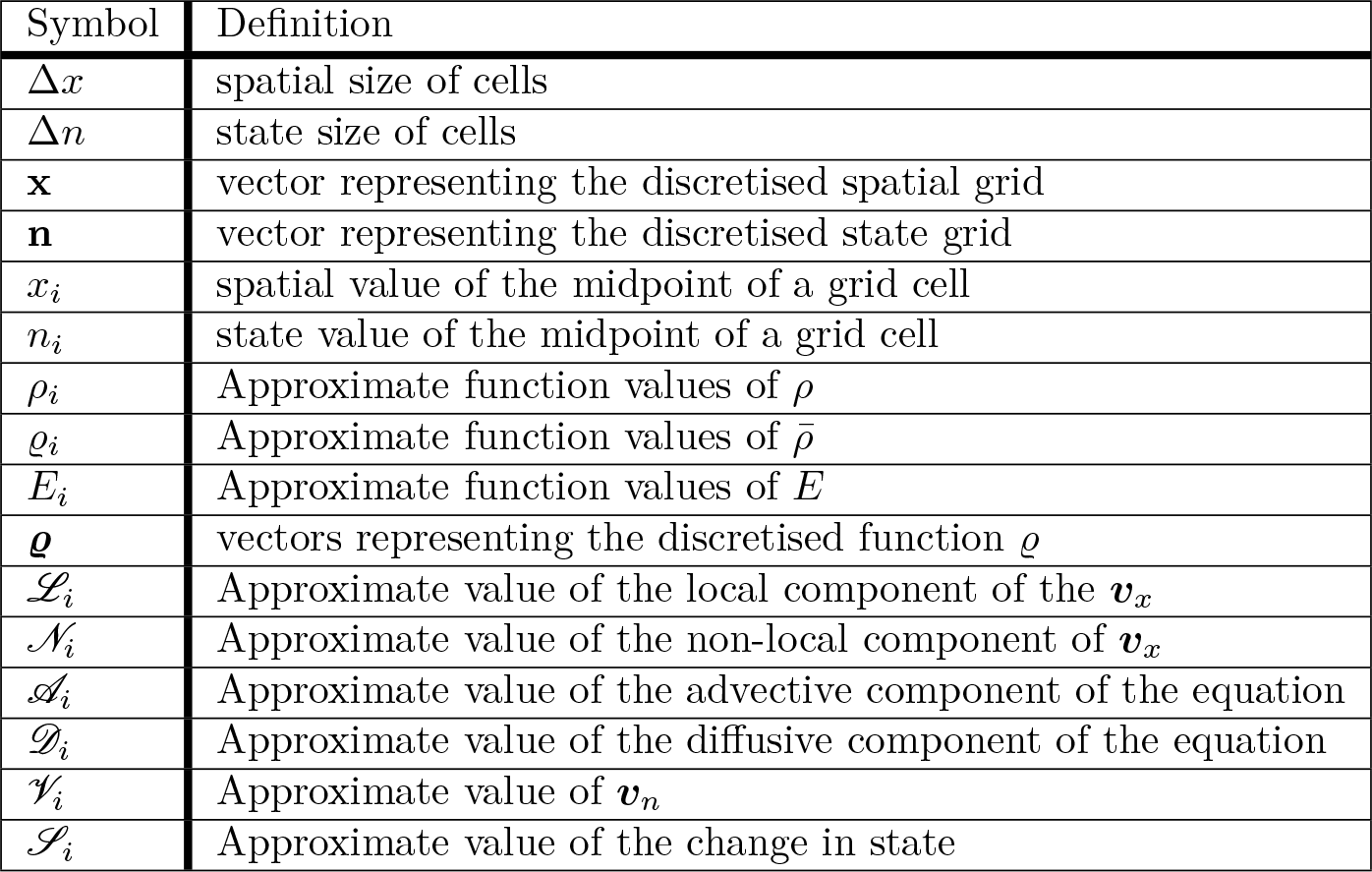
Definitions of symbols used in numerical scheme at arbitrary cell *i* = (*i*_***x***_, *i*_***n***_).

Beginning with the local part of the velocity term (denoted *ℒ*)

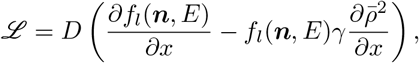

we approximate both derivatives using central differencing schemes, giving

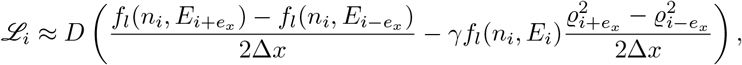

at an arbitrary cell *i*. Then, for the non-local component of the velocity term (denoted *𝒩*),

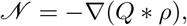

we begin by exploiting the convolution theorem, which states that under suitable conditions the Fourier transform of a convolution of two functions is equal to the point-wise product of their individual Fourier transforms, i.e.,

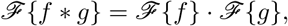

where *ℐ* represents the Fourier transform (we also denote the inverse Fourier transform as *ℐ* ^*−*1^). Additionally, we use the following property of convolutions

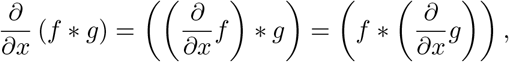

to turn the convolution component of the advection term into

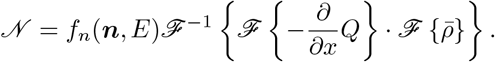

We can then approximate the convolution as

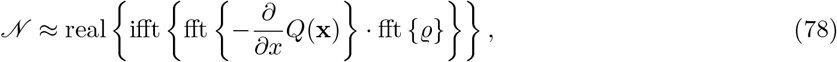

where fft and ifft represent the fast Fourier transform and inverse fast Fourier transform respectively. We take only the real component of the ifft as any imaginary value will simply be due to error. By combining the local and non-local components and letting

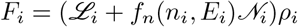

we can approximate the wavespeed at a cell boundary, 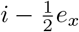, as

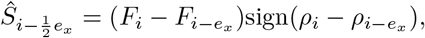

and

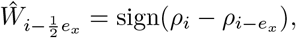

where

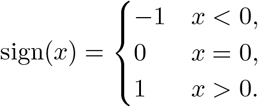

Giving the upwinding scheme for the advection component of movement as

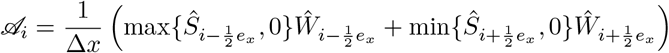

Next, for the diffusion term, *𝒟*

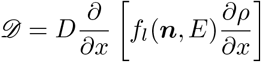

We can approximate this using FVM as

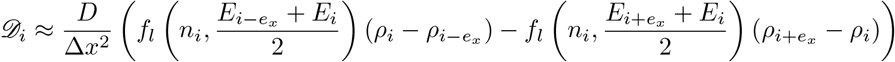

For the state component

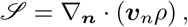

we begin by directly calculating the wave speed at the cell boundaries

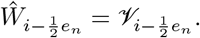

We then find the flux through the cell boundary due to state velocity using the upwinding scheme

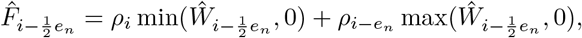

giving

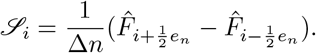

Combining all the terms we obtain,

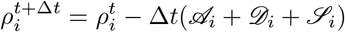

For the CPU case we use an adaptive Dormand-Prince method [19] for the time component and for the GPU case we use an adaptive scheme RK4 based on the work of Horsea and Shampine [26]. Finally all code is available in the following [git repository](https://gitlab.com/FillipeGeorgiou/state-space-numerical-code).

